# Germinal center activity and B cell maturation promote protective antibody responses against *Plasmodium* pre-erythrocytic infection

**DOI:** 10.1101/2021.09.10.459481

**Authors:** Ganesh Ram R. Visweswaran, Kamalakannan Vijayan, Ramyavardhanee Chandrasekaran, Olesya Trakhimets, Samantha L. Whiteside, Vladimir Vigdorovich, Ashton Yang, Andrew Raappana, Alex Watson, William Selman, Meghan Zuck, Nicholas Dambrauskas, Alexis Kaushansky, D. Noah Sather

**Author notes:** Equal contribution (Ganesh Ram R. Visweswaran, Kamalakannan Vijayan).

## Abstract

Blocking *Plasmodium*, the causative agent of malaria, at the asymptomatic pre-erythrocytic stage would abrogate disease pathology and prevent transmission. Rodent-infectious species of *Plasmodium* such as *P. yoelii (Py)* serve as key tools to study vaccine efficacy and disease biology in immune-competent experimental animals. Here we evaluated the differences in vaccine-elicited humoral immunity in two widely used, and vastly diverged, inbred mouse strains, BALB/cJ and C57BL/6J, and identified immunological factors associated with protection. We vaccinated with *Py* circumsporozoite protein (CSP), the major surface antigen on the sporozoite, and evaluated protective efficacy after mosquito bite challenge. Vaccination achieved 60% sterile protection and otherwise delayed blood stage patency in BALB/cJ mice, whereas; all C57BL/6J mice were infected similar to controls. Interestingly, protection was mediated by antibodies, and could be passively transferred from immunized BALB/cJ mice into naïve C57BL/6J. Dissection of the underlying immunological features of protection revealed early deficits in antibody titers and polyclonal avidity in C57BL/6J mice. Additionally, *Py*CSP-vaccination in BALB/cJ induced a significantly higher proportion of antigen-specific B-cells and class-switched memory B-cell (MBCs) populations than in C57BL/6J mice. Strikingly, C57BL/6J mice also had markedly fewer germinal center experienced, CSP-specific class-switched MBCs compared to BALB/cJ mice. Analysis of the IgG γ chain repertoires by next generation sequencing in *Py*CSP-specific memory B-cell repertoires also revealed higher somatic hypermutation rates in BALB/cJ mice than in C57BL/6J mice. These findings indicate that BALB/cJ mice achieved higher levels of B cell maturation in response to vaccination with *Py*CSP, which likely enabled the development of protective antibody responses. Overall, our study indicates that germinal center activity and B cell maturation are key processes in the development of vaccine-elicited protective antibodies against CSP.

## Introduction

Malaria remains a major public health crisis, with more than 229 million cases resulting in more than 409K deaths in 2019, concentrated in sub-Saharan Africa and disproportionately affecting women and children (WHO, 2020). After peaking in 2004, cases have steadily declined, but in recent years cases have plateaued or slightly increased, highlighting the urgent need for new counter measures to achieve eradication (WHO, 2019). Vaccines that prevent infection with *Plasmodium* parasites, the causative agents of malaria, offer the best hope to overcome this plateau and facilitate eradication. While vaccines are in development for all stages of the *Plasmodium* life cycle, the pre-erythrocytic (PE) stage is an attractive target, as stopping the parasite at this asymptomatic stage would prevent infection, subsequent disease, and transmission (Burrows et al., 2017). RTS,S, the only malaria vaccine with regulatory approval (Olotu et al., 2016; Stoute et al., 1997; Sun et al., 2003), and the most clinically advanced whole sporozoite vaccine, *Pf*SPZ (Sissoko et al., 2017), both target this stage and result in only partial protection in field trials. Recently, a new PE vaccine candidate, R21, achieved promising results in early clinical field trials (Datoo et al., 2021). Subunit vaccines, such as RTS,S and R21, induce potent antibody responses against the major surface protein on the sporozoite, the <u>C</u>ircum<u>s</u>porozoite <u>P</u>rotein (CSP) (Datoo et al., 2021; Sun et al., 2003). Whole sporozoite vaccines also elicit anti-CSP antibodies, they also produce antibodies to other *Plasmodium* antigens (Camponovo et al., 2020; Mordmuller et al., 2017).

The mechanisms by which pre-erythrocytic (PE) vaccines prevent malaria infection are yet to be fully characterized, but both antibodies and CD8^+^ T cells have been implicated, depending on the vaccine modality (Epstein et al., 2011; Ewer et al., 2013; Ewer et al., 2015) Antibodies have been shown to mediate anti-parasitic activity and are thought to work primarily in the skin and interstitial tissues where they interfere with the sporozoite’s motility and survival (Aliprandini et al., 2018). This has been confirmed by studies of monoclonal antibodies (mAbs) targeting CSP, which have been shown experimentally to reduce or prevent *Plasmodium falciparum* infection (Aliprandini et al., 2018). The study of anti-CSP mAbs isolated from humans has implicated antibody affinity, epitope specificity, and B cell clonal selection as key factors mediating protective function (Murugan et al., 2018). Together these studies indicate that antibodies are a key mediator of protection in subunit PE vaccines, and that protective antibodies have inherent features that determine neutralizing capacity. Whereas mAbs have been instrumental to our understanding of antibody-mediated protection from malaria, vaccination with CSP induces complex polyclonal responses consisting of a diversity of antibodies. The key features of vaccine-elicited polyclonal antibody responses that determine protection from infection have yet to be fully defined, but their characterization would be a critical milestone toward the development of an effective CSP-based vaccine.

Here, we aimed to identify the characteristics of protective antibody responses elicited by full-length CSP vaccination using the rodent malaria, *P. yoelii*. This model of malaria infection enabled us to conduct live vaccination and mosquito bite challenges in a wild type experimental system to dissect the correlates of antibody-mediated protection. To identify features associated with efficacy, we characterized vaccine-elicited B cell responses in two highly diverged mouse strains, BALB/cJ and C57BL/6J, that exhibited differential vaccine-mediated sterilizing immunity. We evaluated serum antibody responses, characterized CSP-specific B cell phenotypes, and compared B cell receptor repertoires between the two strains in response to CSP vaccination. We found that vaccine-elicited anti-CSP antibodies alone are sufficient to achieve sterile protection from infection, and that protection was associated with higher magnitude germinal center (GC) responses and somatic mutation of CSP-specific B cells. These findings implicate B cell maturation as a critical determinant in the development of potent sterilizing antibody-mediated immunity against malaria and indicate that vaccine modalities aimed at inducing mature B cell responses will be necessary to achieve sterilizing immunity in the field.

## Results

### Immunization with *Py*CSP elicits anti-parasitic antibodies and sterile protection from *P. yoelii* in BALB/cJ, but not C57BL/6J mice

The major strains used to study vaccine efficacy and host pathogen interactions in murine models of malaria infection are BALB/cJ and C57BL/6J (Benhnini et al., 2009; Kuipers et al., 2017). In our studies of CSP vaccine-elicited vaccine efficacy, we observed that these two strains exhibited differential levels of protection in a subunit immunization, mosquito bite challenge model, where we routinely failed to achieve sterilizing immunity in C57BL/6J mice. To compare efficacy directly, we immunized animals (n=10 per group) with 20 μg of full length ecto-domain recombinant *Py*CSP in 20% adjuplex adjuvant at weeks 0, 2 and 6 (Figure 1A). Control groups received PBS-formulated with 20% adjuplex. At weeks 3 and 7, blood samples were collected to evaluate the antibody responses elicited by the vaccination. At week 7, both mouse strains were subjected to *P. yoelii* 17XNL sporozoite infection by mosquito-bite challenge (15 mosquitoes/mouse) and were monitored for three weeks or until blood-stage patency was reached. All the placebo-injected control mice in both the BALB/cJ and C57BL/6J groups developed blood-stage patency in as early as 5 days post mosquito bite challenge, indicating that the strains were equally susceptible to infection (Figure 1B), as has been reported previously (Miller et al., 2014). All of the *Py*CSP-vaccinated C57BL/6J mice were unprotected and developed blood-stage patency 6 days post mosquito bite challenge. Surprisingly, in contrast to the C57BL/6J mice, *Py*CSP-immunized BALB/cJ mice exhibited sterile protection from infection (60%). Six BALB/cJ mice were parasite free at the end of the experiment, and the remaining mice (40%) exhibited a significant delay in blood stage patency (7 days) (Figure 1B and S1A).

**Figure 1.**
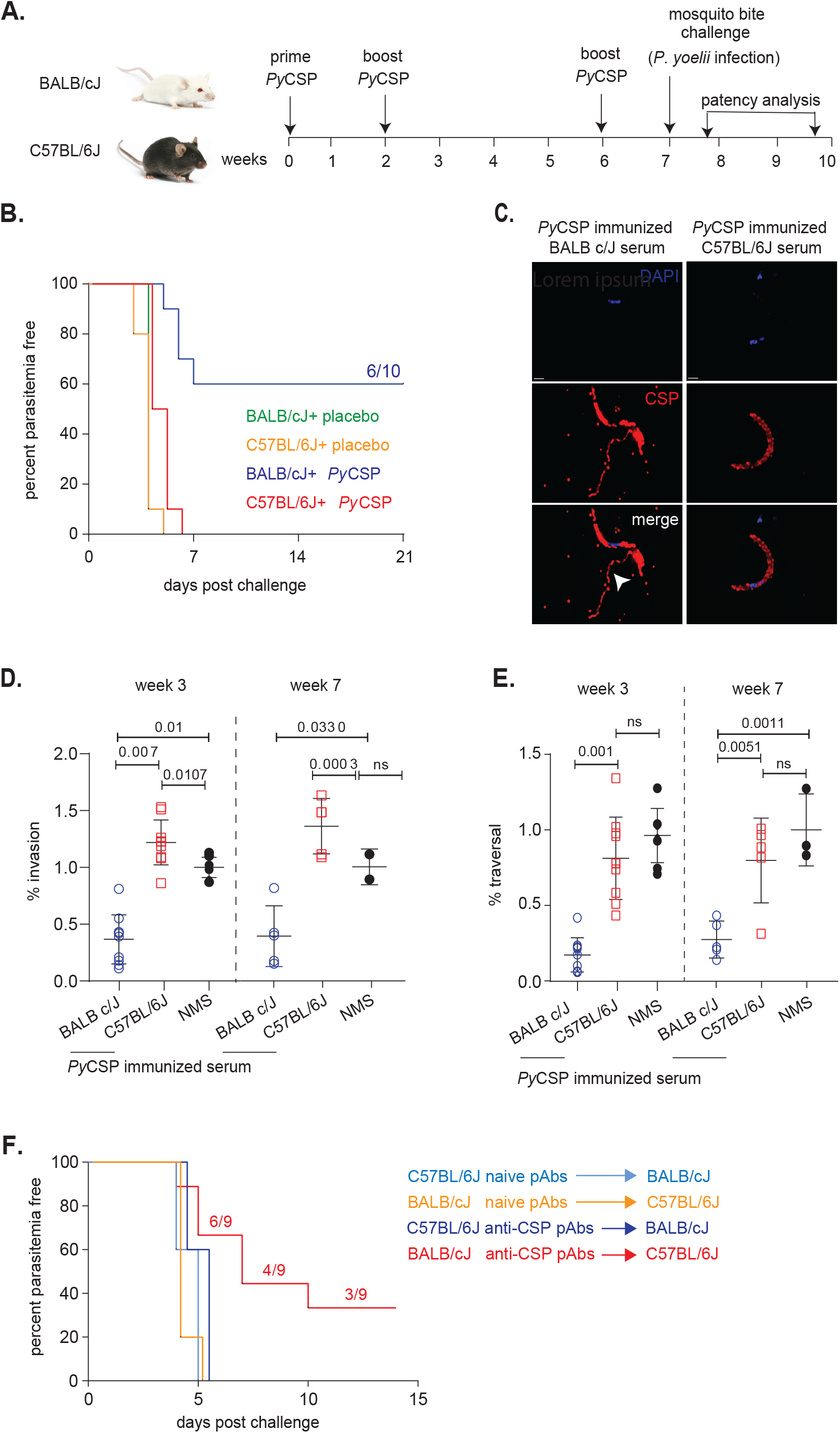
The *Py*CSP-immunization in BALB/cJ and C57BL/6J mice delays blood stage patency and confers protection only in the prior. **A**. The immunization and challenge regimen are illustrated. 6-8 weeks old female BALB/cJ (n=10) and C57BL/6J (n=10) mice were intramuscularly immunized with 20 µg of *Py*CSP formulated with 20% adjuplex on weeks 0, 2 and 6 (arrows) followed by *Py* sporozoites-bearing mosquito bite challenge (15 mosquitoes/mouse) on week 7. Post challenge the blood smears were collected from the tail vein for up to 3 weeks to monitor blood-stage patency by Giemsa stain. Submandibular bleeds were performed in Week 3 and on week 7 (before challenge). **B**. BALB/cJ and C57BL/6J were immunized with *Py*CSP at 0, 2 and 6 weeks. A week post final boost, mice were challenged with *P. yoelii* infected mosquitoes and patency was assessed from day 3 through day 21. The Kaplan-Meier Survival plots represent percentage parasitemia free mice over time. Data are from 10 mice /condition across 2 independent experiments. The control, *Py*CSP-immunized BALB/cJ and C57BL/6J mice are indicated in green, orange, blue and red, respectively. The numeric indicates that 6 out of 10 *PyCSP*-immunized BALB/cJ mice were parasitemia free. **C**. Representative immunofluorescent images of *P. yoelii* sporozoites incubated with purified serum antibodies from post week 7-*Py*CSP-immunized BALB/cJ and C57BL/6J mice for 10 min. *Py*CSP monoclonal antibody, 2F6, is used as control for CSP trailing assay and DAPI is used to stain the nucleus of the sporozoites. The arrowhead points to the regions of CSP reaction or shedding (CSPR). (**D** and **E**) Freshly isolated sporozoites were pre-incubated with *Py*CSP-immunized BALB/cJ and C57BL/6J serum antibodies were incubated with *Py* sporozoites (1:1 dilution) for 20 mins. Normal mouse serum (NMS) is used as infection control for *in vitro* assays. Hepa 1-6 cells were infected with antibody-treated sporozoites for 90 min to assess hepatocyte entry. The bar graph here represents the percentage of hepatocytes that were CSP-positive as evaluated by flow cytometry (**D**). Antibody-treated sporozoites were exposed to HFF-1 cells in the presence of Dextran-FITC for 30 mins. The bar graph represents the percentage of cells traversed as assessed by dextran positive cells **(E)**. For (**D**) and (**E**), data represents mean values ± SE from three independent experiments: n=3. p values were determined by comparing each treatment to untreated using one-way ANOVA for multiple comparisons tests. **F**. Polyclonal antibodies from *Py*CSP-immunized BALB/cJ and C57BL/6J mice sera were purified and *Py*CSP-specific antibodies were quantified by ELISA. Naïve BALB/cJ (n=5) and C57BL/6J (n=9) were passively transferred (i.p.) with 90 µg of *Py*CSP antibodies from *Py*CSP-immunized C57BL/6J (blue) and BALB/cJ (red) mice, respectively. Post five days of antibody passive transfer mice were challenged with mosquito-bites from 15 *Py-*infected mosquitoes and the blood stage patency is monitored for 2 weeks. Naïve BALB/cJ (n=5) and C57BL/6J (n=5) that were passively transferred with purified polyclonal antibodies from naïve C57BL/6J (light blue) and BALB/cJ (orange) mice, respectively were used as controls. Data analyzed by Two-way ANOVA and p values were obtained by Tukey’s multiple comparison test. \*\**\* p<0*.*0004; **p<0*.*005; *p<0*.*0005;* ns- not significant.

To characterize the underlying immune responses mediating sterile protection, we analyzed the anti-parasitic activity of serum polyclonal antibodies (pAbs) *in vitro* and *in vivo*. Anti-CSP serum antibodies from both strains recognized CSP on the surface of the *P. yoelii* sporozoite by immunofluorescence microscopy (Figure 1C), indicating that both strains developed antibodies capable of recognizing surface displayed CSP on the sporozoite. However, only BALB/cJ anti-CSP pAbs induced a CSP reaction on sporozoites (Figure 1C, arrows) in which neutralizing antibodies (Nabs) induce the sporozoite to “shed” its coat of CSP (Balaban et al., 2021; Ghosh and Jacobs-Lorena, 2009; Vanderberg et al., 1969) We then characterized *in vitro* inhibitory activity in the inhibition of Sporozoite Traversal and Invasion (ISTI) assay. *Py*CSP-immunized BALB/cJ and C57BL/6J serum antibodies from Weeks 3 and 7 were pre-incubated with *Py* sporozoites and the percentage invasion and traversal to hepatocytes were analyzed. *Py*CSP-immunized BALB/cJ immune serum from both weeks 3 and 7 significantly inhibited sporozoite invasion into hepatocytes compared to controls, whereas *Py*CSP-immunized C57BL/6J immune serum from weeks 3 and 7 did not inhibit invasion above background levels (Figure 1D). Similarly, *Py*CSP-immunized BALB/cJ pAbs significantly reduced sporozoite traversal at weeks 3 and 7 through hepatocytes compared to controls, whereas *Py*CSP-immunized C57BL/6J immune serum did not inhibit traversal above background levels at any time (Figure 1E).

To evaluate the differential functional activity *in vivo*, and to assess potential infection differences due to mouse strain, we performed a pAb swap infection challenge and sterile protection experiment. BALB/cJ or C57BL/6J mice were immunized with *Py*CSP as shown in Figure 1A and then protein A purified pAbs from immunized (Post 3^rd^ immunization) animals or naïve controls were passively infused into strain mis-matched naïve mice five days prior to mosquito bite challenge (15 mosquitoes/mouse). Mice were monitored for two weeks or until blood-stage patency was reached. Thus, C57BL/6J immune and naïve serum was infused into naïve BALB/cJ mice prior to challenge, and vice versa. The pAbs were purified from pooled serum and the anti-CSP content was quantified using a standard curve generated with canonical neutralizing anti-*Py*CSP mAb, 2F6. Each animal received an equivalent of 90 µg of polyclonal anti-CSP antibody, with naïve animals receiving 90 µg total IgG. C57BL/6J mice that received BALB/cJ-derived anti-CSP pAbs exhibited delays to patency and 40-60% sterile protection, depending on the experiment (Figure 1F and S1B), whereas BALB/cJ mice infused with C57BL/6J-derived anti-CSP pAbs were not protected and all became infected similar to controls (Figure 1F and S1B). These findings demonstrate that the protection observed in vaccinated BALB/cJ mice is mediated by antibodies alone. Further, it implies that there are specific features of vaccine-elicited antibodies in BALB/cJ mice that are critical to achieving sterile protection, features that likely do not exist in C57BL/6J mice. This differential model of sterile protection from malaria afforded the opportunity to assess what characteristics of the B cell response are associated with protection from infection.

### *Py*CSP immunized C57BL/6J mice exhibit early deficits in the development of anti-CSP antibodies

To investigate the underlying differences in the protection between *Py*CSP-immunized BALB/cJ and C57BL/6J mice, we analyzed *Py*CSP-specific plasma IgG responses over the course of vaccination. Antibodies are often not measurable after the first immunization but reach high levels with the anamnestic response after the boost immunizations. Thus, we focused on studying samples from after the second and third immunizations, at weeks 3 and 7, respectively. At week 3, α-CSP IgG endpoint titers were nearly one log_10_ higher in BALB/cJ mice than in C57BL/6J mice (Figure 2A). However, by week 7, one week post third immunization and just before mosquito-bite challenge, IgG titers were similar between the strains (Figure 2A). No difference in IgG subclass usage was observed at week 7, and both strains had similar α-CSP titers of IgG1 and IgG2b antibodies (C57BL/6J do not express Ig2a) (Figure S2). Thus, although we detected an early difference in overall binding titers, at the time of challenge the strains had similar levels and subclass distribution of circulating α-CSP IgG antibodies.

**Figure 2.**
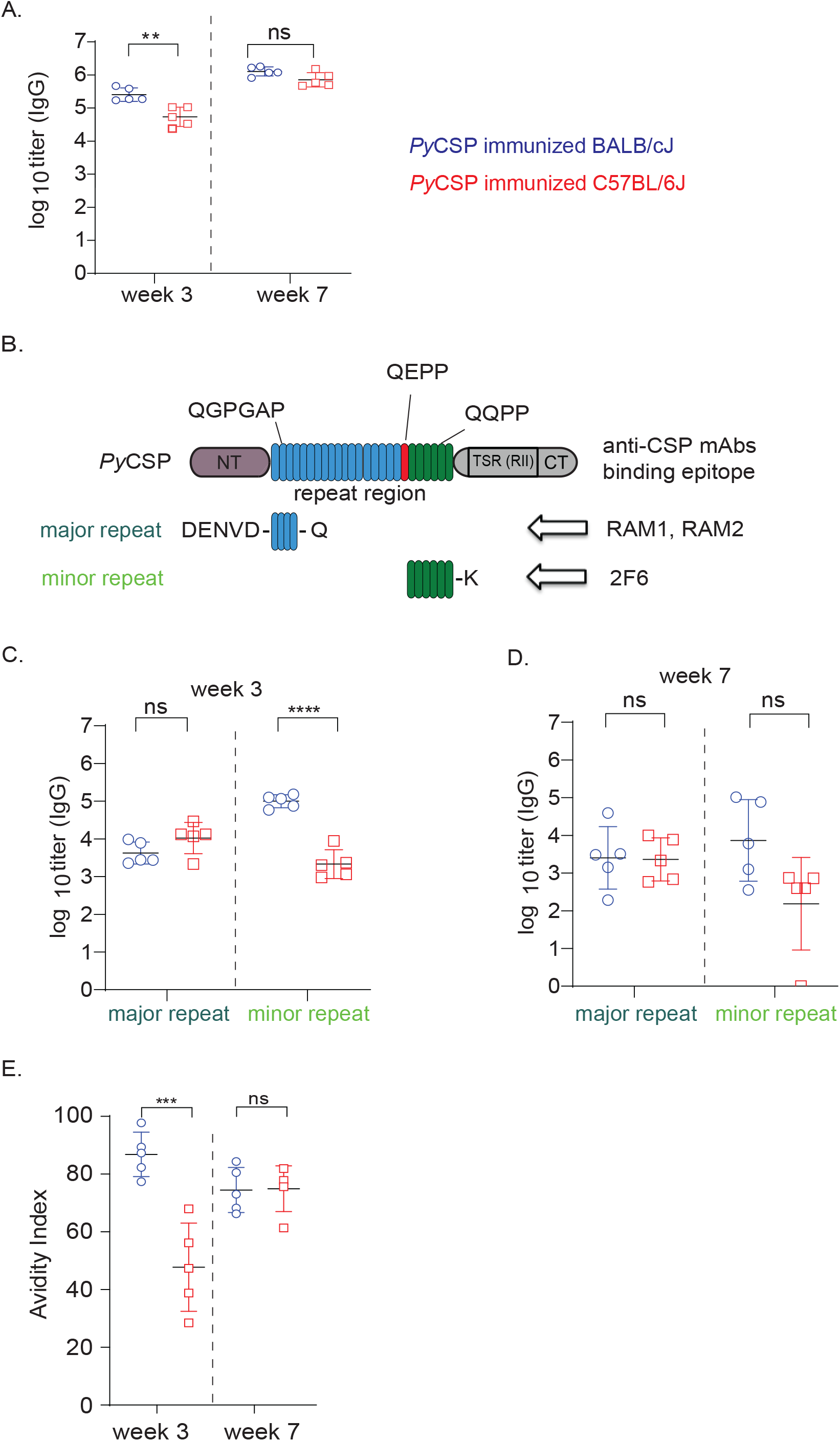
Comparison of antibody responses after immunization of *Py*CSP at week 3 and 7 time points in BALB/cJ and C57BL/6J mice. **A**. 6-8 weeks old female BALB/cJ (n=5) (blue) and C57BL/6J (n=5) (red) mice were intramuscularly immunized with 20 µg of *Py*CSP with 20% adjuplex on weeks 0, 2 and 6 and the blood samples were collected on week 3 and week 7. ELISA was performed to analyze the *Py*CSP-specific total IgG titers. **B**. Cartoon representing the domain organization of mature *Py*CSP ectodomain. The N- and C-terminal domains are represented as NT and CT, respectively. The TSR (RII) domain is in CT. The major and minor repeats of the repeat domain are colored blue and green, respectively and their respective amino acid repeat sequences are illustrated. The QEPP amino acid sequence connecting the major and minor repeat regions is indicated in red. The binding epitopes of anti-*Py*CSP mAbs RAM1, RAM2 and 2F6 are indicated with arrows. **C** and **D**. *Py*CSP-specific IgG titers of *Py*CSP-immunized BALB/cJ (blue) and C57BL/6J (red) mice to the major and minor repeats at Week 3 **(2C)** and Week 7 **(2D)** were analyzed. **E**. The avidity of the *Py*CSP-immunization elicited antibodies to the *Py*CSP antigen at weeks 3 and 7 in BALB/cJ (blue) and C57BL/6J (red) mice were analyzed. Avidity index is calculated by (AUC of NH_4_SCN wells)/ (AUC of PBS wells)*100. Data analyzed by Two-way ANOVA and p values were obtained by Tukey’s multiple comparison test. *\*\*\*\*p<0*.*0001; ***p<0*.*0004 **p<0*.*005;* ns- not significant.

We next measured the avidity index of the serum IgG to full ectodomain PyCSP over the course of immunization, which is a surrogate measure of the strength of polyclonal antibody binding to the protein. Binding is measured in the presence and absence of thiocyanate chaotrope, and the relative disruption of binding is used to generate an index value (Pullen et al., 1986). Interestingly, C57BL/6J mice had significantly lower avidity for recombinant *Py*CSP after the second immunization at week 3 (Figure 2E), indicating a potential early deficit in antibody maturation. By week 7 the avidity indices against full length rec-*Py*CSP were statistically similar. However, this lack of difference in polyclonal avidity at week 7 is likely due to the course measure of the binding characteristics of a diverse mixture of antibodies as a population, where the potential presence of less frequent high affinity clones may not be sufficient to influence the overall avidity index.

To determine whether differential protection could be attributed to epitope specificity, we generated repeat peptides corresponding to the major and minor repeat regions (Figure 2C) and assessed IgG binding over the course of immunization. The repeat regions are known to be the target of neutralizing antibodies, whereas the N- and C-terminal domains have not been implicated as targets for neutralizing antibodies (Kisalu et al., 2018; Scally et al., 2018; Tan et al., 2018; Thai et al., 2020; Triller et al., 2017). The repeat region in *Py*CSP contains two general motifs that make up the longer, N-terminal major repeat region and the shorter, more C-terminal minor repeats (Figure 2B). The minor repeat region is the target of potent α-*Py*CSP neutralizing antibody activity of mAb 2F6 (Bruna-Romero et al., 2004; Sack et al., 2014; Vijayan et al., 2021) but neutralization targeting the major repeat has not been reported. At week 3, the strains had equivalent IgG titers to the major repeat peptide, but BALB/cJ mice had significantly higher titers to the minor repeat peptide (Figure 2D). At week 7, IgG titers to the major repeat remained statistically similar, but, although the difference was not statistically significant, C57BL/6J titers to the minor repeat trended lower (Figure 2D). These findings imply that the unprotected phenotype in C57BL/6J mice may be due to dampened antibody responses to the minor repeat, which is known to mediate neutralization.

### Both major and minor repeat motifs of *Py*CSP mediate sterile protection from *P. yoelii* infection

To evaluate this possibility, we investigated whether the major repeat also could be a target for NAbs, or whether neutralization is solely mediated through the minor repeat motif as observed for mAb 2F6. We previously characterized a non-neutralizing antibody (nNAb), RAM1, which binds to the major repeat with relatively low affinity, does not induce the CSP reaction, nor protects *in vitro* or *in vivo* (Vijayan et al., 2021). In this study, we isolated a mAb RAM2, which also binds to an epitope in the major repeat motif like RAM1 (Figure 3A and 3D). RAM2 bound to recombinant CSP with a similar EC_50_ to that of 2F6 by ELISA (Figure 3B), and both mAbs had similar binding kinetics measured by Octet-BLI (Figure 3C). In contrast to RAM1, RAM2 induces the CSP reaction (Figure 3E) and blocks *in vitro* infection in ISTI to similar levels the NAb 2F6 (Figure 3F). Unlike RAM1, RAM2 also facilitates sterile protection *in vivo* when administered by passive infusion at survival levels that mirror 2F6 (Figure 3G), indicating that potent neutralization can target either the major or minor repeat motifs, and that the differential protection between mouse strains cannot be fully explained by differential responses to the major and minor repeat motifs.

**Figure 3.**
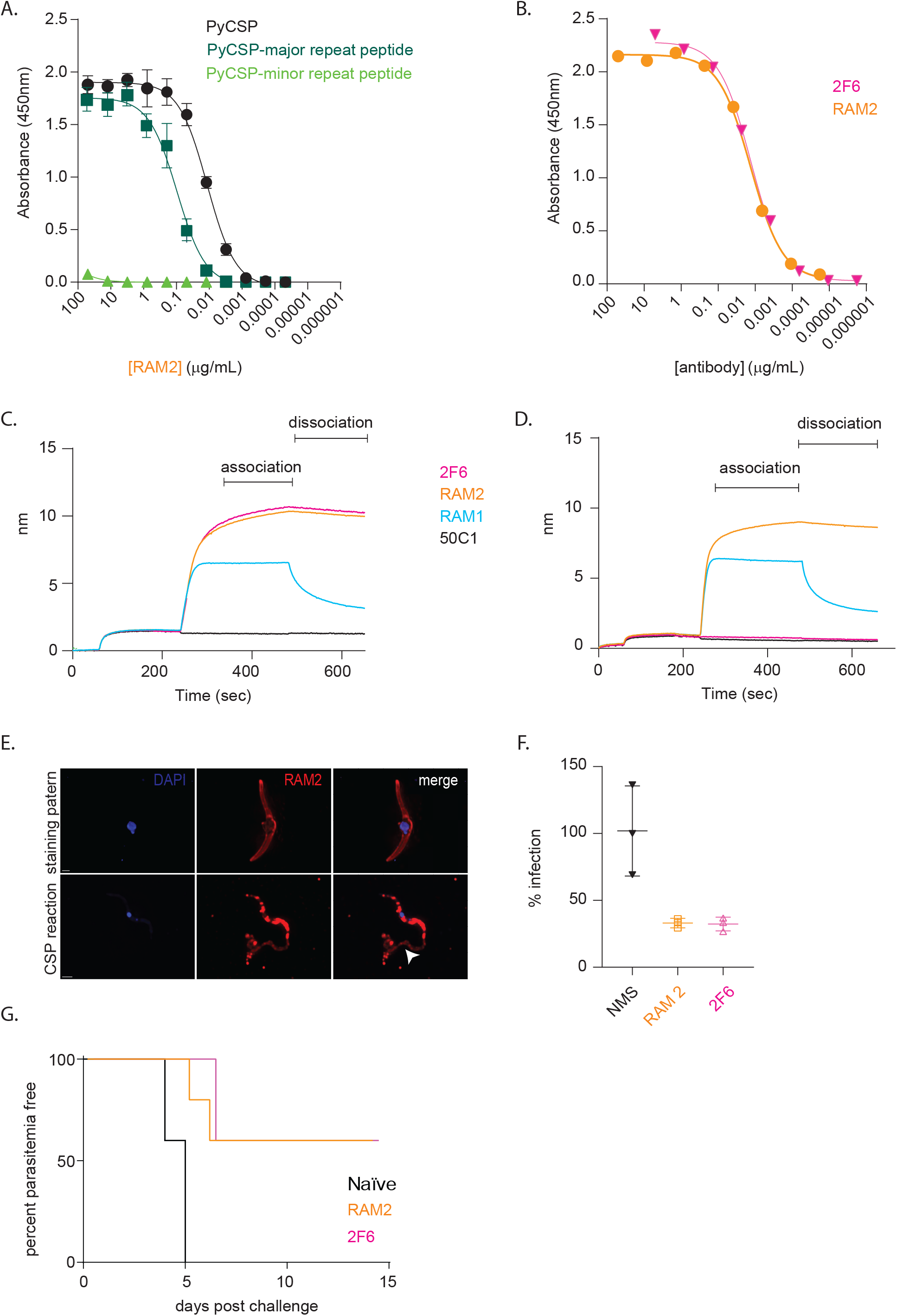
*Py*CSP mAb RAM2 avidly binds to major repeat regions of *Py*CSP and provides sterile liver stage protection from the pathogen. **A**. ELISA showing the binding of RAM2 to the *Py*CSP full length protein (black), the major repeat peptide (dark green) but not to the minor repeat peptide (light green). **B**. ELISA showing the EC_50_ of RAM2 (orange) and 2F6 (pink) binding to *Py*CSP full length protein **C**. Biotinylated-*Py*CSP (5 µg) derivatized streptavidin biosensors were incubated in different mAbs (RAM1 (cyan), RAM2 (orange) 2F6 (pink) and 50C1 (black), 5 µg each) and the association and dissociation kinetics were assayed by Octet-BLI **D**. Biotinylated-major repeat peptide (5 µg) loaded streptavidin biosensors were dipped in 5 µg each of mAbs (RAM1-cyan, RAM2-orange, 2F6-pink and 50C1-black) and the association and dissociation kinetics were assayed by Octet-BLI. **E**. Representative immunofluorescent images of *P. yoelii* sporozoites incubated with mAb RAM2 (10 μg) for 10 min. *Py*CSP mAb, 2F6, is used as control for CSP trailing assay and DAPI is used to stain the nucleus of the sporozoites. The arrowhead points to the regions of CSP reaction or shedding (CSPR). **F**. Freshly isolated sporozoites were pre-incubated with (10 µg) of RAM2 (orange), 2F6 (pink) and normal mouse serum (black) antibodies for 10 mins. Hepa 1-6 cells were infected with antibody-treated sporozoites for 90 min to assess hepatocyte entry. The bar graph here represents the percentage of hepatocytes that were CSP-positive as evaluated by flow cytometry. The data represents mean values ± SE from three independent experiments: n=3. p values were determined by comparing each treatment to untreated using one-way ANOVA for multiple comparisons tests. **G**. BALB/cJ were injected with 150 µg of RAM2 (orange) or 2F6 (pink) intraperitoneally. Naïve BALB/cJ (black) mice injected with PBS is used as control. After 24 h, mice were challenged with bites from fifteen *P. yoelii* infected mosquitoes and patency was assessed from day 4 through day 14. Kaplan-Meier survival plot represents percentage parasitemia free mice over time, including 10 mice from 2 independent experiments.

Although RAM1 and RAM2 both target the major repeat domain, they do so with drastically different binding kinetics. RAM2 Fab binds to recombinant CSP with a K_D_ 7.85 × 10^−7^ M and with an association and dissociation kinetics values of K_on_ (1.25 × 10^4^ 1/Ms) and K_off_ (9.84 × 10^−3^ 1/s), whereas RAM1 Fab exhibits little binding to CSP and we were unable to derive a measurable K_D_ (Figure S3; Vijayan et al., 2021). By comparison, the minor repeat NAb 2F6 Fab binds to rec-*Py*CSP with a K_D_ of 5.24 × 10^−7^ M (Vijayan et al., 2021). This difference in binding affinity between RAM1 and RAM2 is likely a major contributor to their differential functional activity and implies that there may be a minimum threshold for affinity that is necessary to achieve neutralization. Further, these findings raise the possibility that the lack of neutralization in C57BL/6J mice may be due to a lack of high affinity B cell clones, potentially driven by lower B cell maturation. Taken together, these findings implicate antibody affinity as a potential key feature of protective antibody responses against *Plasmodium*.

### Dampened B cell responses elicited by vaccination in C57BL/6J mice

We next evaluated if differential antibody protection between BALB/cJ and C57BL/6J mice was a consequence of nuanced responses to *Py*CSP vaccination in the B cell compartment. To assess the potential immunological origins of the differential protection, we characterized anti-CSP-specific B cell responses in the spleen after vaccination. Splenocytes were harvested at week 3 and 7 or post 1 week after the second and third immunizations, respectively, and stained with markers for flow cytometric analyses, including B220, GL-7, CD38, IgD, IgM, and CD138. Recombinant CSP was tetramerized by conjugation to streptavidin-conjugated APC and APC/Fire750, which were used for column enrichment and dual staining for *Py*CSP-reactive cells. We assessed and quantified the total number of splenic *Py*CSP-specific B-cells (CD3^-^, B220^+^, CSP^+^), IgM memory B-cells (MBCs) (CD3^-^, B220^+^, CD38^+,^ CD138^-^, IgM^+^, IgD^-^, CSP^+^), germinal center (GC) experienced MBCs (CD3^-^, B220^+^, GL7^+^, CSP^+^), and class-switched MBCs (CD3^-^, B220^+^, CD38^+,^ CD138^-^, IgM^-^, IgD^-^, CSP^+^) (Figure 4 and S4). Both immunized animals and strain-matched placebo immunized animals were analyzed.

**Figure 4.**
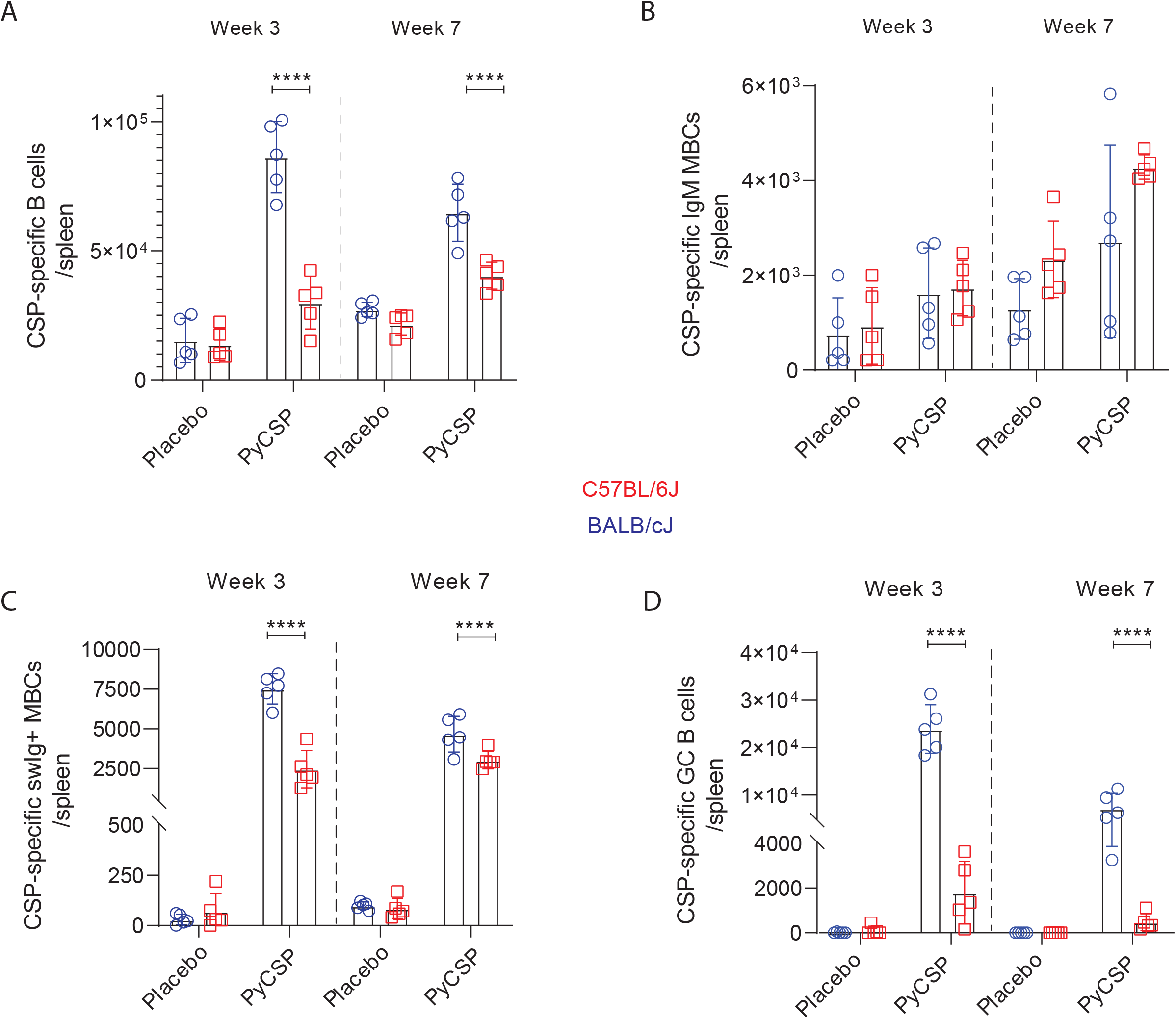
*Py*CSP-specific B-cell responses of BALB/cJ and C57BL/6J mice at weeks 3 and 7. 6-8 weeks old female BALB/cJ (n=5) and C57BL/6J (n=5) mice were immunized intramuscularly with 20 µg of *Py*CSP adjuvanted with 20% adjuplex on weeks 0, 2 (Week 3) or weeks 0, 2 and 6 (week 7) and the splenocytes were isolated either on week 3 or week 7 as described in the materials and methods. Cartoon representing the general workflow in B-cell population analysis is illustrated (Figure S4). B-cells responses of different B-cell populations per spleen were quantified for *Py*CSP-specific B-cells **(A)**, IgM memory B-cells (MBCs) **(B)**, class-switched (sw) Ig+MBCs **(C)**, and Germinal center (GC) B-cells **(D)**. Data analyzed by Two-way ANOVA and p values were obtained by Tukey’s multiple comparison test. *\*\*\*\*p<0*.*0001*.

Interestingly, *Py*CSP-immunized BALB/cJ mice exhibited significantly higher numbers of splenic *Py*CSP-specific B-cells after the 2^nd^ and 3^rd^ immunizations (Figure 4A), indicating a more robust anamnestic response than in vaccinated C57BL/6J mice. This difference was especially apparent in week three, which follows the second immunization, where CSP-specific B cells were nearly 3-fold higher. We did not detect differences in the frequency of CSP^+^ IgM MBCs after either immunization (Figure 4B), although their frequency was relatively low in all samples. However, *Py*CSP-immunized BALB/cJ had more than 3-fold higher numbers of class-switched CSP^+^ MBCs at week three after the second immunization and statistically higher numbers at week 7 (Figure 4C). Strikingly, BALB/cJ mice had approximately 10-fold higher numbers of splenic GL7^+^, GC-experienced CSP-specific B cells than C57BL/6J mice after the second immunization (week 3), and approximately 5-fold more after the third immunization (week 7) (Figure 4D). Thus, except for IgM^+^ MBCs, BALB/cJ mice exhibited comparatively higher *Py*CSP-specific B cell responses, GC activity, and class switch in response to vaccination. As these timepoints measure the anamnestic response, these findings also indicate marked differences in recall responses between the strains. The relative paucity of GC-experienced and class switched MBCs in C57BL/6J mice is intriguing, and likely explains the early differences in IgG titers and binding avidity observed after the second immunization. Overall, the relative deficit GC activity and class switched MBCs may contribute to the inability of vaccinated C57BL/6J mice to produce antibodies that can mediate sterile protection from infection.

### *Py*CSP-immunized BALB/cJ mice achieve higher levels of somatic mutation in response to vaccination

It is possible that the differences in germinal center activity affect overall maturation and somatic mutation within the CSP-specific B cell receptor (BCR) repertoire, which ultimately drives the generation of high affinity antibodies after vaccination. To assess this, we sorted class-switched CSP-reactive MBCs after either two or three immunizations with recombinant PyCSP and sequenced their IgH heavy chain B cell receptor repertoire by 5’RACE next generation sequencing. Class-switched CSP-specific MBCs (CD3^-^, B220^+^, IgM^-^ IgD^-^, CD38^+^, anti-CSP^+^) were stained and enriched as described above, and then sorted into RLT lysis buffer by fluorescent activated cell sorting (FACS). Class switched MBCs from unimmunized animals were also sorted to serve as a reference for sequence analysis. The sorted, CSP-specific MBCs in mice within each strain were combined, and then IgG transcripts were amplified by 5’ Rapid amplification of cDNA ends (5’RACE) and sequenced on an Illumina MiSeq instrument, allowing the retrieval of the entire V, D, J rearrangement. The resulting sequences were processed in our in-house bioinformatics pipeline for processing and V/D/J gene annotation and were analyzed to determine the percent identity of each BCR from germline alleles (i.e., rates of somatic mutation) and third complementarity determining region of antibody heavy chains (CDRH3) characteristics. Higher percent identity to germline indicates less mutation, and conversely, lower identity indicates higher rates of mutation. We annotated and estimated the percent sequence identity to V-region germline of the CSP-specific BCR repertoire at weeks three and seven (i.e., one week post two or three immunizations, respectively), as well as the total class switched MBC IgG BCR repertoire from unimmunized animals (Figure 5). The CDRH3 size distributions of the sequenced γ chain repertoires are shown in Figure S5.

**Figure 5.**
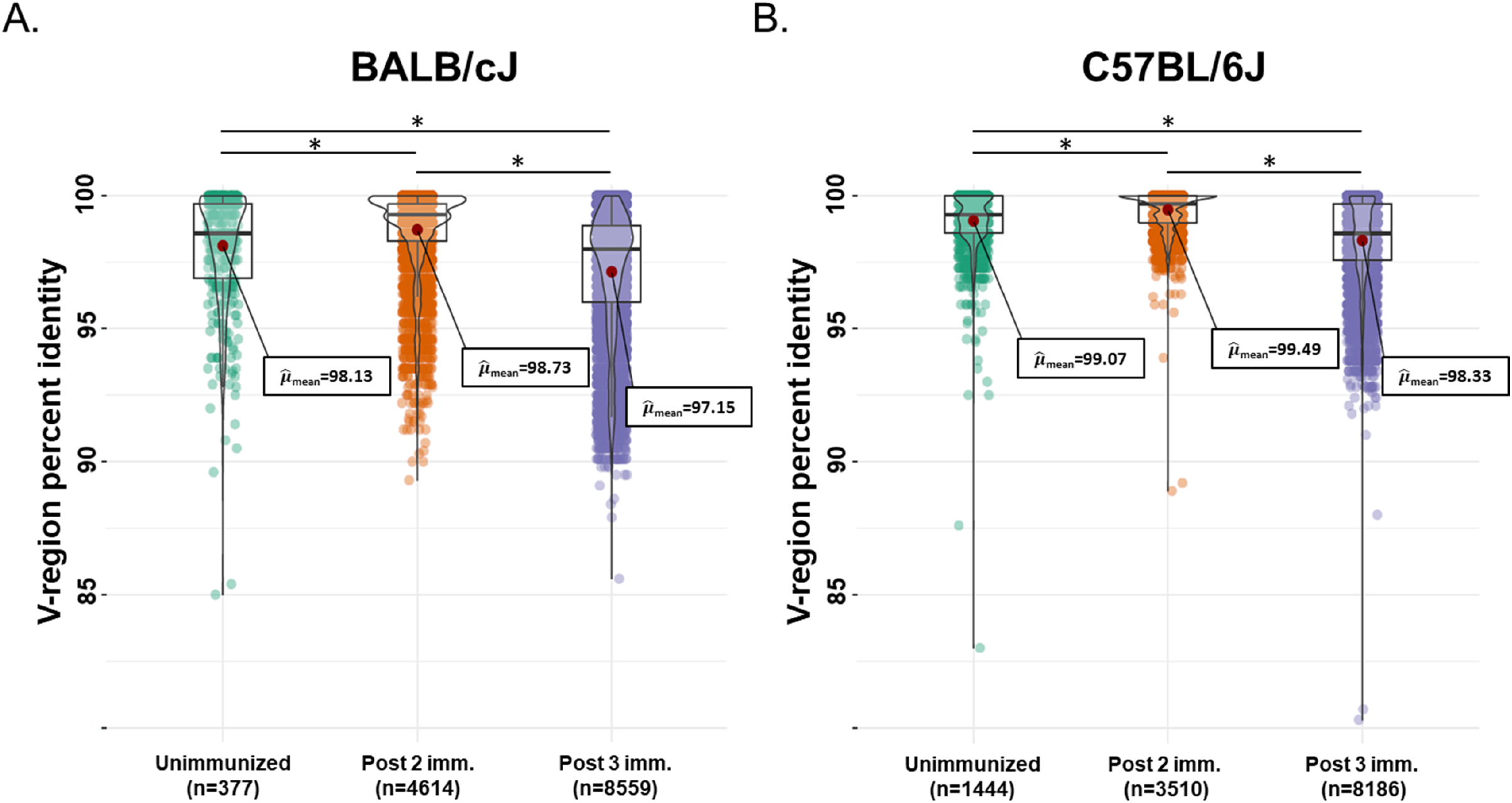
*Py*CSP-immunized BALB/cJ mice employ more highly diverse V-gene rearrangements than C57BL/6J. Somatic hypermutations in the V-gene of IgG heavy chains of *P*yCSP-specific mouse memory B-cells at Week 3 (orange, post 2^nd^) and 7 (purple, post 3^rd^) performed by high throughput cell sorting and NGS in BALB/cJ **(A)** and C57BL/6J **(B)** mice. The unimmunized mice (green) in a similar age group of the week 7 vaccinated mice were used as controls. The results are representative of three independent experiments (n=5/group). Each data point represents a sequence containing deduplicated members (i.e., 100% identity) and supported by at least 10 raw reads. The visualizations were analyzed and generated using the ggstatsplot (version 0.7.2) R package.

The average CSP-specific MBC V-region identity after two immunizations in BALB/cJ mice was 98.73%. After three immunizations, identity dropped significantly to an average of 97.15%, perhaps an indicator of higher levels of affinity maturation and somatic mutation stimulated by the third immunization (Figure 5A). BCR sequences after three immunizations were detected with as low as ∼86% identity, and numerous sequences were identified with less than 92.5% identity. Thus, IgG heavy chains with 7.5-14% somatic mutation were generated in response to vaccination, although these were in the minority in the overall repertoire. The identity to germline of CSP-specific MBCs was statistically higher after two immunizations than the average identity of MBCs in unimmunized BALB/cJ mice (98.73% vs. 98.18%), but lower after three immunizations (98.18% vs 97.15%). Thus, on average, compared to the total MBC repertoire in unimmunized animals, CSP-specific MBCs had less mutation after two immunizations, a possible consequence of BCR clonal selection, but significantly more mutation after three immunizations, suggesting ongoing diversification and affinity maturation.

In contrast, the average identity of CSP-specific IgG heavy chains in C57BL/6J mice was 99.49% after two immunizations and 98.33% after three (Figure 5B), with only very rare outliers detected with more than 7.5% mutation. Thus, the CSP-specific IgG heavy chain repertoire in C57BL/6J mice after two immunizations remained close to germline identity, possibly related to the lack of GC experienced CSP-specific MBCs detected at this timepoint (Figure 4D). Of note, the non-CSP MBC BCR repertoire in unimmunized mice was more diverse in BALB/cJ mice than in C57BL/6J mice (98.13% vs 99.07), which may be indicative of intrinsic underlying differences between the strains. Taken together, these findings indicate that BALB/cJ mice appear to generate higher levels of somatic mutation in response to vaccination with *Py*CSP. It follows that the higher level of mutation more commonly generated in BALB/cJ mice could be a key factor in their ability to generate high affinity anti-CSP antibodies and achieve sterile protection from infection.

## Discussion

Defining the features of antibodies that mediate protection from pre-erythrocytic malaria infection is a key milestone that will guide future vaccine development efforts. As RTS,S rolls out more widely, and newer protein-based vaccines are developed, such as R21, it is essential to understand the key characteristics of vaccine-elicited antibody responses that mediate protection from liver stage infection. Here we studied vaccine responses in two strains of mice, BALB/cJ and C57BL/6J, that differ in their sterile protection from infection in response to vaccination with recombinant *Py*CSP. We studied several aspects of the B cell response to vaccination and identified key differences that implicate B cell maturation as a defining factor in the development of protective antibody responses. These findings have direct implications for vaccine development, highlighting the urgent need for vaccine formulations and regimens that drive potent, mature antibody responses.

Previous studies have established that binding affinity to CSP is a key factor in the ability of monoclonal antibodies to neutralize infection (Epstein et al., 2011; Jongo et al., 2018; Sissoko et al., 2017; Vijayan et al., 2021). High affinity antibodies are generated through the process of affinity maturation. This occurs in the primary and secondary lymphoid tissues, where antigen-reactive B cells proliferate and migrate into germinal centers. Once in germinal centers, they undergo rounds of somatic hypermutation and clonal selection, ultimately resulting in high affinity B cell clones that secrete antibodies. The effect of affinity on neutralization is exemplified by the mAbs RAM1 and RAM2. RAM2 is protective, whereas RAM1 is not, despite binding to the same peptide repeats. In this scenario, affinity is the major factor driving the difference in neutralization; RAM2 binds with an affinity several orders of magnitude higher than RAM1.

Thus, the significant lack of germinal center experienced B cells in C57BL/6J mice is striking, as it indicates that this process is either impaired or happening far less frequently than in BALB/cJ mice. These observations dovetail with our finding that less somatic hypermutation from germline occurred in C57BL/6J mice within the CSP-specific class-switched MBC compartment, indicating that less B cell maturation occurred after vaccination. In fact, the CSP-specific BCR repertoire population average remained close to germline identity, even after the anamnestic response of the second immunization. The natural consequence of less B cell maturation is likely that fewer, less diverse high affinity BCR clones are generated after vaccination, or that overall, the population may be less mutated, as we observed. This is also evident in the lower polyclonal avidity after the second immunization, which may be a significant underlying factor in the lack of antibody protection. Each of these observations support the model where less efficient germinal center activity and class switch (and, in turn, insufficient high-affinity CSP antibodies) in C57BL/6J mice leads to a lack of sterilizing protection, whereas robust affinity maturation activity in BALB/cJ mice, resulting in a subset of high-affinity NAbs like RAM2 and 2F6, leads to sterilizing protection (Figure 6).

The underlying immunological origins of reduced germinal center activity in C57BL/6J mice remain unclear and could relate to potential differences in HLA or MHC alleles or subpar T helper responses. However, vaccine protection from *P. berghei* rodent malaria infection has been reported in C57BL/6J mice, especially with adjuvants that strongly drive B cell development, such as Matrix M (Collins et al., 2021). Further, in this study passive transfer of CSP-specific antibodies elicited in BALB/cJ mice achieved sterile protection in C57BL/6J mice. This indicates that it is possible to elicit or passively transfer antibodies in C57BL/6J mice that mediate sterile protection. As such, it is likely that our vaccine regimen was not sufficient to elicit antibodies with the desired characteristics in C57BL/6J, despite eliciting very high titers of anti-*Py*CSP antibodies. Therefore, it is not that C57BL/6J mice were generally hypo-responsive to vaccination or overly sensitive to infection, but rather a deficit in maturing the B response toward ideal neutralization characteristics likely underlies the total lack of protection. Thus, although we do not know the underpinnings of the differential antibody responses in C57/BL6J mice, comparative use of this model allowed the identification of fundamental immunological processes that could be key parameters for achieving durable sterilizing immunity.

It is intriguing that C57BL/6J mice lacked any measurable protection from infection, despite statistically equivalent polyclonal antibody avidity indices in the samples taken just prior to mosquito bite challenge (Figure 2E). The avidity index is based on chaotrope disruption of binding in ELISA and is a course measure of the binding behavior of antibodies as a population mixture (Alexander et al., 2015; Pullen et al., 1986). This measure is extensively used as a surrogate measure of aggregate B cell maturation. However, this measure can be influenced by nuances in epitope specificity and valency, and may under measure more rare high affinity, highly functional antibodies (Alexander et al., 2015). Much of the antibody response may not be relevant or effective for sterile protection, and it is likely that only a subset of B cell clones will produce antibodies with the desired characteristics. Given the equivalent IgG titers and avidity indices just prior to infection challenge, it is possible that that the protection in BALB/cJ mice is mediated by relatively rare, highly mutated, high affinity B cell clones, rather than through a large population of antibodies that may be less mutated and lower affinity. To deconvolve this will likely require further examination at single B cell level, where direct affinity/functional relationships can be determined, as we have done for RAM1 and RAM2. Indeed, in studies of natural infection and whole sporozoite vaccination, only a tiny fraction of antibodies obtained are functional against the parasite, implying that potent functional antibodies are rare in the overall B cell repertoire (Camponovo et al., 2020).

Eliciting durable, potent antibody responses by vaccination is a considerable challenge for the field. RTS,S, which contains the potent adjuvant AS01, elicits high titers of CSP antibodies, but titers and partial protection wanes quickly after vaccination (Datoo et al., 2021). R21, which is similar to RTS,S, is currently in testing with the potent adjuvant Matrix M, which is hoped to be superior in eliciting high titers of antibody responses (Datoo et al., 2021). However, our findings imply that high titers of antibodies alone are not sufficient to achieve potent, durable protection from infection. Rather, it is the quality of the antibodies in the response that will determine whether protection can be achieved. A careful examination of B cell responses, including the extent of their hypermutation and germinal center experience, may be critical for extracting meaningful correlates of protection and, ultimately, designing the most potent vaccine regimen against malaria. Novel approaches that emphasize the elicitation of high affinity antibodies to neutralizing epitopes will likely be a more fruitful path forward for vaccine development. Such a vaccine would dramatically impact our ability to curtail malaria infection worldwide and could make a substantial impact in the push towards eradication.

## Materials and Methods

### Cloning and production of *Py*CSP and truncation constructs

*P. yoelii* CSP, a 403-amino acid (aa) protein, consisting of an N- and C-terminal domains connected by a central repeat domain (Figure 2B). The two predicted *N*-glycosylation sites (S27A, T348A), one each in the N- and C-terminal domains of the native antigen were mutated to prevent the addition of N-linked glycans in the mammalian expression system and the non-glycosylated protein, *Py*CSP (38.3 kDa), is used in this study. The cartoon representing the *Py*CSP, and different major and minor repeat peptides used in this work are illustrated in Figure 2B and the construction and expression of the protein is published elsewhere(Vijayan et al., 2021). The *Py*CSP antigen was codon optimized for human bias and C-terminally 8X His and Avi-tagged, connected via a GS-linker to the antigen, to facilitate purification and biotinylation. A tPA signal was added at the N-terminus to facilitate protein secretion and the construct was cloned into pcDNA3.4 vector (Thermo Fisher, Waltham, MA, USA) which drives transcription via a CMV promoter. After sequence confirmation the plasmid DNA (500 ng/ml of cells) encoding the antigens were introduced into HEK293 suspension cultures (1 million/ml) by high-density transfection using 2 mg/ml polyethyleneimine (PEI) (1:5 Pei:plasmid DNA). Cells were grown in FreeStyle 293 serum-free media (Thermo Fisher) for five days at 37°C 8% CO_2_. Cells were spun down by centrifugation (4000 rpm for 20 min at 4°C) and the supernatant was collected. Sodium azide was added to a final concentration of 0.02% and NaCl was added to a final concentration of 500 mM. The antigens were purified by a two-step chromatography protocol. The proteins were captured using Ni-NTA (Qiagen, Germantown, MD, USA), washed in EQ buffer (25 mM Tris pH 8.0, 300 mM NaCl, 0.02% NaN3), and eluted in EL buffer (25 mM Tris pH 8.0, 300 mM NaCl, 200 mM Imidazole, 0.02% NaN3). The elution fractions were collected, pooled, and concentrated and further purified on a standardized Superdex 200 16/600 (GE Healthcare, Chicago, IL, USA) column running in HBS-E (10 mM HEPES, pH 7, 150 mM NaCl, 2 mM EDTA). Peak fractions were pooled, concentrated, and stored at 4°C until use.

### Animal immunization

All animal studies were conducted under protocols reviewed and approved by the Institutional Animal Care and Use Committee (IACUC) at Seattle Children’s Research Institute. For regular immunization of 6-8 weeks old female BALB/cJ and C57BL/6J mice, 20 µg of *Py*CSP formulated in 20% adjuplex is administered intramuscularly at weeks 0, 2 and 6. Blood samples were collected at one week post 2^nd^ and 3^rd^ immunizations i.e., week 3 and week 7, by submandibular or chin bleeds. Placebo-injected mice were used as experimental control. For studying the B-cell responses and NGS of the antigen-specific memory B-cells, mice were harvested at week 7 followed by splenocyte isolation. For pathogen challenge experiments, all the mice were exposed to *P. yoelii* infected mosquito bite challenge (10-15 mosquitoes/mouse) in week 7 and followed for 2-3 weeks for blood-stage patency. For serum antibodies swap passive transfer experiment, all the BALB/cJ mice were injected i.p., with 90 µg *Py*CSP-antibodies purified from *Py*CSP-immunized C57BL/6J mice and vice versa followed by mosquito bite challenge (10-15 mosquitoes/mouse) 5 days post passive transfer of antibodies. As a control, purified naïve BALB/cJ serum antibodies (90 µg) were i.p., transferred to C57BL/6J mice and vice versa. All the mice were monitored for 2-3 weeks for blood stage patency.

### ELISA

Plasma antibody binding to *Py*CSP and to different truncation variants and peptides were determined using a Streptavidin-capture ELISA. Mouse plasma samples were heat-inactivated for 30 min at 56°C prior to all assays. All ELISA incubations were done for 1 h at 37°C and plates were washed between each ELISA step with PBS, 0.2% Tween-20. To determine antibody binding to the ligand, Immulon 2HB 96-well plates (Thermo Scientific, 3455) were coated with 50 ng/well of Streptavidin (NEB, N7021S) in 0.1 M NaHCO_3_, pH 9.5, followed by blocking with 3% BSA in PBS. Later the plates were coated with 200 ng/well with biotinylated ligand antigen or peptides followed by a blocking step 10% non-fat milk, 0.3% Tween-20 in PBS. Later mouse plasma was serially diluted in duplicate over a range of 1:200 to 1:55,987,200 in PBS with 0.2% BSA. *Py*CSP mAb, 2F6, was serially diluted on each plate to ensure intra-assay consistency. Bound antibodies were detected using goat anti-mouse IgG Fc-HRP (Southern Biotech) or goat anti-mouse IgG1 Fc-HRP (Southern Biotech) or goat anti-mouse IgG2a Fc-HRP (Southern Biotech) or goat anti-mouse IgG2b Fc-HRP (Southern Biotech) at 1:2000 dilution in PBS with 0.2% BSA. Plates were developed with 50 μl/well of TMB Peroxidase Substrate (SeraCare Life Sciences Inc, 5120-0083) and stopped after 3 min with 50 μl/well of 1 N H_2_SO_4_. Absorbance at 450 nm was read using a BioTek ELx800 microplate reader. Binding curves were generated by nonlinear regression (log[agonist] vs response [three parameters]) in GraphPad Prism V8 (San Diego, CA). Endpoint titers were defined as the reciprocal of plasma dilution at OD 0.1.

### Chaotrope Dissociation ELISA

Antibody avidity of post-2^nd^ and 3^rd^ (Week 3 and week 7) immunization plasma samples to *Py*CSP were determined using direct immobilization ELISA. Immulon 2HB 96-well plates (Thermo Scientific, 3455) were coated with 50 ng/well of *Py*CSP overnight at room temperature, then blocked the following day with PBS, 10% non-fat milk, 0.3% Tween-20 for 1 h at 37°C. Plates were washed between each step with PBS-containing 0.2% Tween-20. Mouse plasma samples were heat-inactivated for 30 min at 56°C prior to all assays. Plasma was diluted in PBS, 10% non-fat milk, 0.03% Tween-20 over a range of 1:50 to 1:9,331,200 and plated in quadruplicate side-by-side on the same plate. After a 1 h incubation with antibodies, plates were washed, and half of the sample wells were treated with 2 M NH_4_SCN in PBS, while the other half was treated with PBS alone for 30 min at room temperature. Plates were then washed and goat anti-mouse IgG Fc-HRP (Southern Biotech, 1013-05) was added to the plate at 1:2000 in PBS, 10% non-fat milk, 0.03% Tween-20. Plates were developed as described above. The avidity index was calculated as the ratio of AUC of samples treated with 2 M NH_4_SCN over AUC of samples treated with PBS: (AUC_NH4SCN_ / AUC_PBS_)×100.

### Monoclonal antibodies isolation and production

The method of generation of *Py*CSP monoclonal antibodies were described earlier (Carbonetti et al., 2017; Vijayan et al 2021). Briefly, six weeks old female BALB/cJ mice were immunized with 20 µg of *Py*CSP adjuvanted with 20% v/v of Adjuplex at 0, 2 and 6 weeks followed by spleens harvest and splenocyte isolation at Week 7. Using EasySep mouse B cell enrichment kit and manufacturer’s protocol (StemCell Technologies Inc., Tukwila, WA, USA), B cells were isolated by negative selection and resuspended in FACS buffer (PBS with 2% Fetal calf serum) followed by staining with anti-mouse CD16/CD32 (mouse Fc block; BD Biosciences) and a decoy tetramer (BV510) for 10 mins at room temperature. Later the cells were stained with CD38-APC, IgM-FITC, IgD-AF700, B220-PacBlue (BioLegend, San Diego, CA, USA) and *Py*CSP tetramer (BV786) for 30 mins at 4°C followed by a wash with FACS buffer. Finally, the cells were resuspended in FACS buffer and filtered by using a 30-micron filter followed by single-cell sorting using a BD FACS Aria II with a 100-μm nozzle running at 20 psi. *Py*CSP-specific class switched memory B cells (B220+ CD38+ IgM-IgD-antigen+ decoy-cells) were isolated and sorted as single cell per well into 96-well PCR plates. For the cDNA generation and the consequent IgG and IgK variable region amplification a previously described (von Boehmer et al., 2016) protocol was followed and the PCR amplified final heavy and light chain sequences were cloned by Gibson assembly into pcDNA3.4 expression vectors containing the murine IgG1 and IgK constant regions, respectively. The IgG and IgK plasmid DNA sequences were verified for the CDR3 sequences by Sanger sequencing and these plasmid pairs were diluted in PBS followed by addition of polyethyleneimine (Polysciences, Warrington, PA). After 15 mins of incubation at room temperature, the mixture was added to HEK293-F cells at a density of 1 million cells per milliliter cultured for 5 days in mammalian FreeStyle™ at 37°C, 5% CO2 on a shaker platform for mAb expression. The cells were separated from the supernatant by centrifugation at 4000 rpm for 20 mins at room temperature and the cell-free supernatant was passed over a HiTRAP MabSelect Sure column (GE Lifesciences #11003493) followed by wash and elution steps performed as suggested by the manufacturer. The isolated mAbs were buffer exchanged into HBS-E (10 mM HEPES, pH 7, 2 mM EDTA, 150 mM NaCl) and the homogeneity and the size of the antibodies were analyzed by using an analytical Superdex 200 10/300 column (GE Healthcare, Chicago, IL, USA).

### Tetramer production

The following tetramers were made for antigen enrichment of splenocytes at an 8:1 ratio of protein to SA-fluorophore, based on protein biotinylation efficiency. Bio-*Py*CSP protein (38.3 kDa) at 16 μM was combined with SA-APC antibody (Biolegend) at 2 μM. Bio-*Py*S23 protein (44.8 kDa) at 16 μM was combined with SA-APC/Fire750 antibody (Biolegend) at 2 µM and was used as a decoy. To achieve optimal binding, an antibody was added to protein in two additions with 20 min incubations at room temperature following each addition.

### Splenocyte isolation and Antigen-specific cell enrichment

Splenocytes were isolated from fresh spleens by passing the tissue through a 70 μm strainer and rinsing with splenocyte buffer (1X phosphate buffered saline (PBS) supplemented with 2% fetal bovine serum (FBS), 100 μg/mL DNase I (Sigma) and 5 mM MgCl_2_. Collected cells were spun down at 350 x g for 10 min and resuspended in the FACS buffer (1X PBS supplemented with 2% FBS). Fc block (BD Biosciences) was added to each sample at a 1:100 dilution with decoy tetramer (Bio-*Py*S23, SA-APC/Fire750) and samples were incubated for 10 min at RT. Then the *Py*CSP tetramer (Bio-*Py*CSP, SA-APC) was added, and samples were incubated for 30 min at 4°C. After the incubation, cells were washed with FACS buffer and spun at 300x g for 10 min at 4°C. Anti-APC microbeads (Miltenyi Biotec) in FACS buffer were added and samples were incubated for 15 min at 4°C. After washing, magnetic separation of cells was performed with LS columns (Miltenyi Biotec) on a quadroMACS separator (Miltenyi Biotec). Cell suspensions were applied to a pre-separation 30 μM filter on separate LS columns. Untetramerized cells without magnetic beads that passed through the column were collected as flow through. Then the columns were removed from the quadroMACS magnet and magnetically labeled cells were flushed out with the FACS buffer and collected. Both the labeled and flow through fractions were spun at 300xg for 10 min and resuspended in the FACS buffer.

### Antibody staining and flow cytometry

The following antibody mixture was made and used to stain the flow through and labeled fractions: B220-BV510 (Biolegend) at 1:20, CD38-FITC (Biolegend) at 1:100, IgM-BV786 (BD Biosciences) at 1:20, IgD-PercpCy5.5 (Biolegend) at 1:80, CD3-APC/Fire (Biolegend) at 1:40, GL7-e450 (ThermoFisher) at 1:40, and CD138-BV605 (BD Biosciences) at 1:80. Samples were incubated on ice for 30 min, washed with FACS buffer and resuspended in cold 1% paraformaldehyde (PFA) in PBS and stored overnight at 4°C until collection of events on the cytometer the following day. For compensation controls, cryopreserved splenocytes were removed from liquid nitrogen storage, quickly thawed at 37°C in water bath for 1 min, washed in RPMI media with DNase (RPMI 1640, Corning +10% FBS + 1:100 Pen/strep + 2 mM L-glutamine +100 µg/mL DNase I + 5 mM MgCl_2_), passed through a 70 μM cell strainer, washed again and finally resuspended in FACS buffer. Single stains controls were made with the following antibodies: B220-BV510 (Biolegend) at 1:20, CD38-FITC (Biolegend) at 1:100, IgM-BV786 (BD Biosciences) at 1:20, IgD-PercpCy5.5 (Biolegend) at 1:80, CD3-APC/Fire (Biolegend) at 1:40, B220-APC (Biolegend) at 1:50, B220-e450 (ThermoFisher) at 1:33, and B220-BV605 (Biolegend) at 1:20. Samples were incubated for 25 min at 4°C then washed with the FACS buffer and resuspended in cold 1% PFA in PBS and stored at 4°C overnight. In the flowcytometry experiments to evaluate the number of specific B-cell populations. 1% PFA was added to a small portion of unstained, labeled and flow-through cells and reserved for cell count analysis. These samples were incubated overnight at 4°C followed by the addition of Accucheck beads (ThermoFisher) the following day and the events were collected on the cytometer. For quantification of specific B-cells an LSR II flow cytometer (BD Biosciences) was used, 1.5-2.5 million events were collected for each sample at a threshold of 15,000. A total of 50,000 events were collected for cell count analysis. All the data were analyzed using FlowJo software (FlowJo LLC).

### Biolayer interferometry (BLI)

Biolayer interferometry (BLI) is used to test the antigen, antibody interactions using an Octet QK^e^ instrument (ForteBio Inc., Menlo Park, CA, USA). Streptavidin (SA) biosensors (ForteBio Inc.,) were used to immobilize the biotinylated antigen or peptide, followed by incubation in 10X Kinetics Buffer (PBS+0.1% BSA, 0.02% Tween-20 and 0.05% sodium azide). The antigen-derivatized probes were then dipped in indicated concentrations of antibodies in 10X Kinetics Buffer, respectively, followed by an incubation step in 10X Kinetics buffer to test the dissociation of the interactions. The resulting sensograms from the association and dissociation phases were normalized to the buffer values and analyzed by a global fit simple 1:1 binding model using the ForteBio data analysis software (version 7.0.1.5). The K_D_ was determined from the estimated on- and off-rates of the samples.

### Fab generation

The RAM2 Fabs were generated by papain digestion and purified using Immobilized papain agarose resin (Thermofisher Scientific) following manufacturer’s protocol. Briefly 1 mg of RAM2 mAb in 400 µl of sample buffer (20 mM Sodium phosphate pH 7.0, 10 mM EDTA) is mixed with 400 µl immobilized papain resin in digestion buffer (20 mM Sodium phosphate pH 7.0, 10 mM EDTA, 20 mM Cysteine.HCl) and incubated overnight on a rotatory shaker at 37°C. Later 1.5 ml of Tris-HCl, pH 7.5 is added to the mixture followed by centrifugation at 2000 x g for one minute at room temperature. The supernatant containing Fabs is separated from the immobilized agarose resin and is passed through a 0.22 µM spin column. Finally, the Fab fragments were isolated by negative selection on a HiTrap MabSelect SuRe cartridge with 10 column volumes of PBS, pH 7.4 and concentrated on spin column to desired concentration.

### Generation of infected *Anopheles stephensi* mosquitoes

*P. yoelii* wild type strain 17XNL (BEI resources) was maintained in Swiss Webster (Envigo, SW) mice. Swiss Webster mice were injected intraperitoneal (i.p.) with 250 µL of infected blood at 3-5% parasitemia. Gametocyte exflagellation rate was checked 4 days post-injection. Mice were then anesthetized with 150 µL Ketamine Xylazine solution (12.5 mg/mL ketamine, 1.25 mg/mL xylazine in PBS) and naïve female *Anopheles stephensi* mosquitoes (3-7 days old) were allowed to feed on them. Mosquitos were maintained at 23°C, 80% humidity with a 12L-12D light cycle. On day ten, the midguts from ten mosquitos were dissected and the number of oocysts was counted as a measure of infection. On day 15 post-infection, the salivary glands of the mosquitoes were dissected to extract sporozoites.

### Mosquito Bite Challenge

Mice were anesthetized by IP injection of 150 µL Ketamine Xylazine solution (12.5 mg/mL ketamine, 1.25 mg/mL xylazine in PBS). Once anesthetized pairs of mice were each placed on a carton of 15 infectious mosquitoes and mosquitoes were allowed to bite through the mesh top for 10 min. Every 30 sec mice were rotated among mosquito cartons to ensure equal exposure of mice and to maximize mosquito probing thereby sporozoite transfer. Mice received subcutaneous PBS injections and recovered from anesthesia under a heat lamp. Mice were checked for the presence of blood-stage parasites (patency) beginning four days post-challenge.

### Assessment of blood stage patency

Patency was checked daily by blood smear 4 days post-infection by tail snip. Slides were fixed in methanol, dried, and then stained with giemsa (1:5 in H_2_O) for 10 min and viewed at 100X oil immersion on a compound microscope and twenty fields of view examined for each smear. Mice were considered patent if 2 or more ring stage parasites were observed.

### Sporozoite quantification

To quantify the number of sporozoites per mosquito, 10-12 mosquitoes’ salivary glands were hand dissected. These glands were then ground with a pestle and spun at 800 rpm to pellet debris. Sporozoites were counted on a hemocytometer.

### Invasion and traversal assay

Freshly isolated sporozoites were exposed to 2F6, RAM2, RAM1 or serum antibodies from *Py*CSP immunized BALB/cJ and C57BL/6J mice for 10 mins at indicated concentrations. 5×10^5^ Hepa1-6 cells were seeded in each well of a 24-well plate (Corning) and infected with antibody exposed *P. yoelii* sporozoites at a multiplicity of infection (MOI)=0.25 for 90 mins. Cells were co-incubated with high molecular mass Dextran-FITC (70 kDa) (Sigma). After 90 mins of infection, cells were harvested with accutase (Life Technologies) and fixed with Cytoperm/Cytofix (BD Biosciences). Cells were blocked with Perm/Wash (BD Biosciences) + 2% BSA for 1 h at room temperature then stained overnight at 4°C with antibodies to CSP-Alexafluor 488 conjugate. The cells were then washed and resuspended in PBS-containing 5 mM EDTA. Infection rate was measured by flow cytometry (Douglass et al., 2015) on an LSRII (Becton-Dickinson) and analyzed by FlowJo (Tree Star).

### Immunofluorescence

Freshly isolated sporozoites were exposed to 2F6, RAM2, RAM1 or serum antibodies from *Py*CSP immunized BALB/cJ and C57BL/6J mice for 10 mins at indicated concentrations. Sporozoites were spun at 17000*g* for 5 mins and washed with 1xPBS-EDTA and fixed with 3.7% PFA for 20 mins. Sporozoites were transferred to 8-well chambered slides and the images were acquired with a 100×1.4 NA objective (Olympus) on a DeltaVision Elite High-Resolution Microscope (GE Healthcare Life Sciences). The sides of each pixel represent 64.5×64.5 nm and z-stacks were acquired at 300 nm intervals. Approximately 5-15 slices were acquired per image stack. For deconvolution, the 3D data sets were processed to remove noise and reassign blur by an iterative Classic Maximum Likelihood Estimation widefield algorithm provided by Huygens Professional Software (Scientific Volume Imaging, The Netherlands).

### Next-generation sequencing (NGS)

CSP-specific MBCs were sorted into RLT buffer supplemented with 2-mercaptoethanol and then IgG gamma chain was amplified by 5’ Rapid amplification of cDNA ends (5’RACE) followed by sequencing on an Illumina MiSeq instrument generating the entire V, D, J rearrangements (Doepker et al., 2020; Simonich et al., 2019; Vigdorovich et al., 2016). Briefly, cell lysate was homogenized using QIAshredder columns (Qiagen, #79654). RNA was extracted from homogenized lysate containing a range of 3×10^3^ to 1×10^5^ cells per sample using a ll Prep DNA/RNA Mini kit (Qiagen, #80204). RNA was purified and concentrated using RNAClean XP beads (Beckman Coulter, #A63987). The concentration and quality of the RNA was determined by an Agilent 2100 Bioanalyzer with the Agilent RNA 6000 Pico Kit (Agilent Technologies, #5067-1513). The oligonucleotides used for the NGS were tabulated (Table 1). Up to 1 µg of RNA was mixed with a gamma chain reverse primer (vv-534-TGCATTTGAACTCCTTGCC) and incubated at 72°C for 3 min to denature the RNA, then cooled to 42°C to anneal the synthesis primer. cDNA was generated by mixing 5x First-Strand Buffer, DTT (20 mM), dNTP mix (ThermoFisher, #18427-013, 10 mM), a template switch adaptor with UMI tag (vv-877-AAGCAGUGGTAUCAACGCAGAGNNNUNNNNUNNNNUNNNNUCTTrGrGrG), Recombinant RNase Inhibitor (Takara Bio/Clontech, #2313A, 40 U/ul), and SMARTScribe RT (Takara Bio/Clontech, #639536, 100 U/ul) with the denatured RNA and incubating for 90 min at 42°C then heating to 70°C for 10 min. Uracil-DNA glycosylase (NEB, #M0280S, 5 U/ul) was added and the reaction was incubated for 1 h at 37°C. RNAClean XP beads were used to purify and concentrate the reaction. A polymerase chain reaction (PCR) was performed to generate variable libraries using cDNA with primers vv-869-AAGCAGTGGTATCAACGCAG, vv-870-KKACAGTCACTGAGCTGCT, vv-872-TACAGTCACCAAGCTGCT, and Q5 2x Master Mix (NEB, #M0492S). The reaction was incubated for 30 sec at 98°C, then cycled 18 times at 98°C for 10 sec, 63°C for 30 sec, and 72°C for 30 sec, and finally incubated at 72°C for 5 min. A cleanup step was performed with SPRI beads (Beckman Coulter, #B23318). The second PCR was performed with primers vv873-CACTCTATCCGACAAGCAGTGGTATCAACG, vv874-GGGCCAGTGGATAGAC, vv876-GGGACCAAGGGATAGAC and Q5 Master Mix. The reaction was incubated using the same conditions as the first PCR, but with an annealing temperature of 60°C and only 8-15 amplification cycles. Another clean up step was performed with SPRI beads. After the second PCR, the NEB Library Prep protocol of the manufacturer was followed and samples were adaptor ligated using the NEBNext End Prep kit (NEB, #E7645). The adaptor-ligated DNA fragments were indexed with the NEBNext Multiplex Oligos for Illumina kit (NEB, E7335). Adapted and indexed libraries were quantified using a KAPA Library Quantification Kit (Roche, KK4873) on a QuantStudio3 Real Time PCR System (ThermoFisher, #A28567). Quantified libraries were pooled to a concentration of 4 nM and denatured following the MiSeq Guide (2). Libraries were denatured in 0.2 N NaOH and diluted to 20 pM. As a sequencing control, PhiX Control V3 (Illumina, #FC-110-3001) was denatured and diluted to 20 pM and was added to the pooled library sample at a 1% spike in. The pooled library sample plus PhiX was sequenced using the MiSeq Reagent Kit v3 (Illumina, #MS-102-3003) on the Illumina MiSeq system.

**Table 1.**
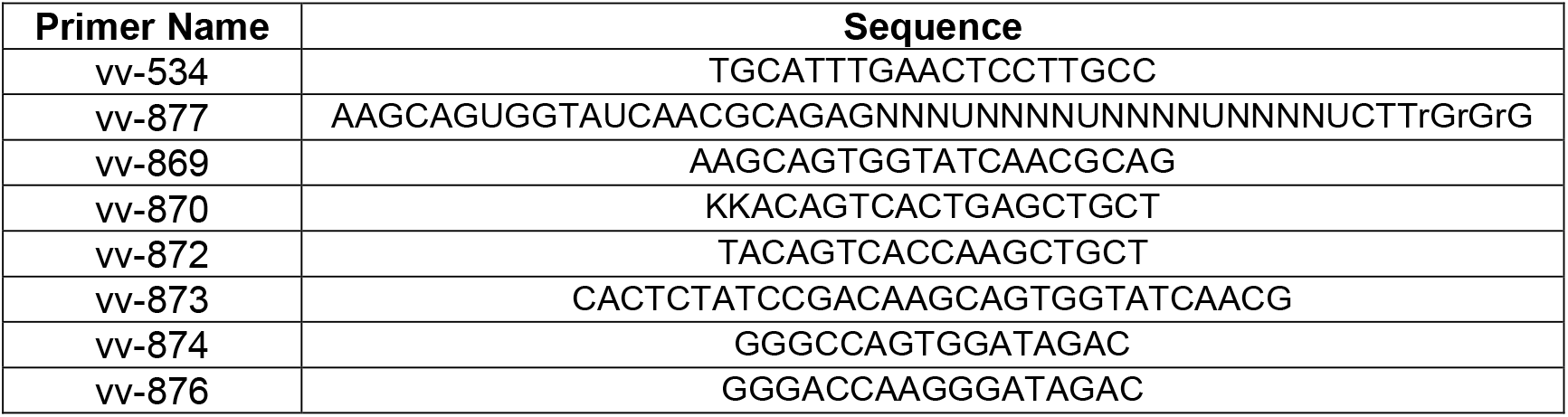
Oligonucleotides used in the NGS experiment.

Raw Illumina MySeq reads were processed using an approach similar to the one previously described (Vigdorovich et al., 2016) with several modifications to utilize unique molecular identifier (UMI)-based error correction (Turchaninova et al., 2016). Briefly, following the amplicon reconstruction and oligonucleotide trimming, UMI sequences were identified and used to collect sequences into molecular identifier groups (MIGs), representing PCR-amplified mRNA molecules. Consensus sequence for each MIG was then determined using the approach adapted from the MIGEC pipeline (Shugay et al., 2014), in which the MIG is first represented by a position frequency matrix, followed by base calling and calculation of a cumulative quality score for each position. Resulting sequence sets were annotated using IgBLAST (version 1.11.0) (Ye et al., 2013) against a custom database of mouse germline sequences obtained from the IMGT/GENE-DB collection (Giudicelli et al., 2005) (www.imgt.org) to determine segment boundaries (e.g., to define CDR3 regions), identify closest germline matches and derive sequence-identity-to-germline values. In order to eliminate multiple identical transcripts likely originating from the same B cell, sequence set deduplication was carried out using VSEARCH (Rognes et al., 2016) (version 2.9.1) at 100% sequence identity. In the finalized deduplicated data sets, only sequences that were supported by ≥10 raw reads were used in further analysis. The visualizations were generated using the ggstatsplot (version 0.7.2) R package.

## Acknowledgments

We thank the vivarium staff at the Seattle Children’s Research Institute for their support and ongoing care of research animals. The authors gratefully acknowledge Rossana de la Noval for administrative and logistical support of this work. This study was funded by a Medical Research Grant from the W.M. Keck Family Foundation to A.K and D.N.S.

## Author Contributions

GRRV, KV, RC, OT, SLW, VV, AY, AR, AW, WS, MZ, and ND conducted experiments and analyzed data. RC, GRRV, KV, MZ, AK, and DNS developed the experimental plan and designed experiments. GRRV, KV, AK, and DNS wrote the manuscript, and all authors edited initial drafts. AK and DNS conceived of the study and obtained funding.

## Figure Legends

**Figure S1.**
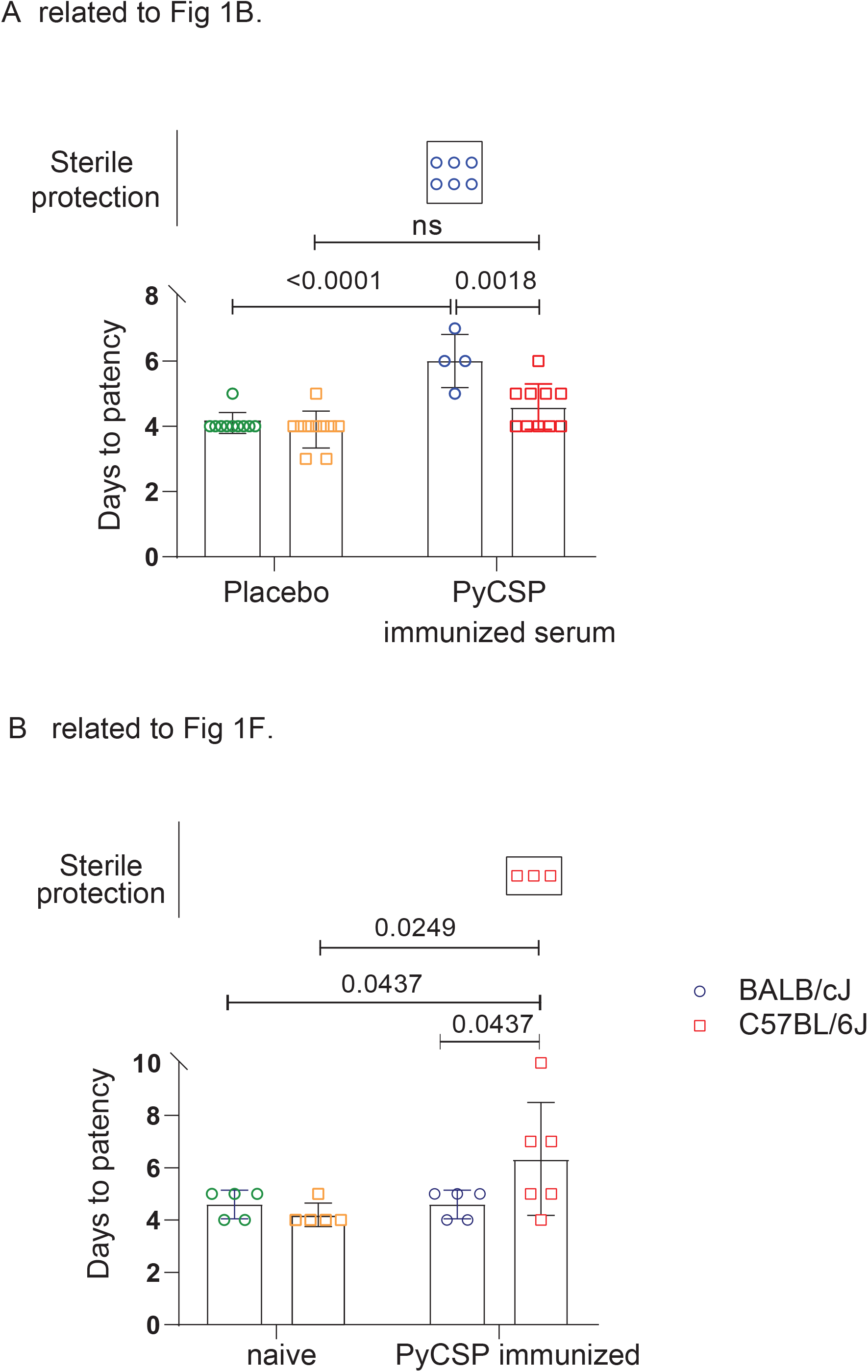
**A**. Graphics showing the statistical significance of *Py*CSP-immunized BALB/cJ (blue), C57BL/6J (red) and their respective placebo controls in green and yellow. **B**. Statistics showing the significance of the delay in blood stage patency in swapped *Py*CSP-pAbs (90 µg)-infused BALB/cJ (blue), C57BL/6J (red) and their respective placebo controls in green and yellow. The days to patency are measured and the number of mice that are sterile protected in the *Py*CSP-immunized BALB/cJ mice **(A)** and *Py*CSP-immunized BALB/cJ mice pAbs infused naïve C57BL/6J mice **(B)** were indicated. Data analyzed by Two-way ANOVA and p values were obtained by Tukey’s multiple comparison test. \*\**\* p<0*.*0004; **p<0*.*005; *p<0*.*0005;* ns- not significant

**Figure S2.**
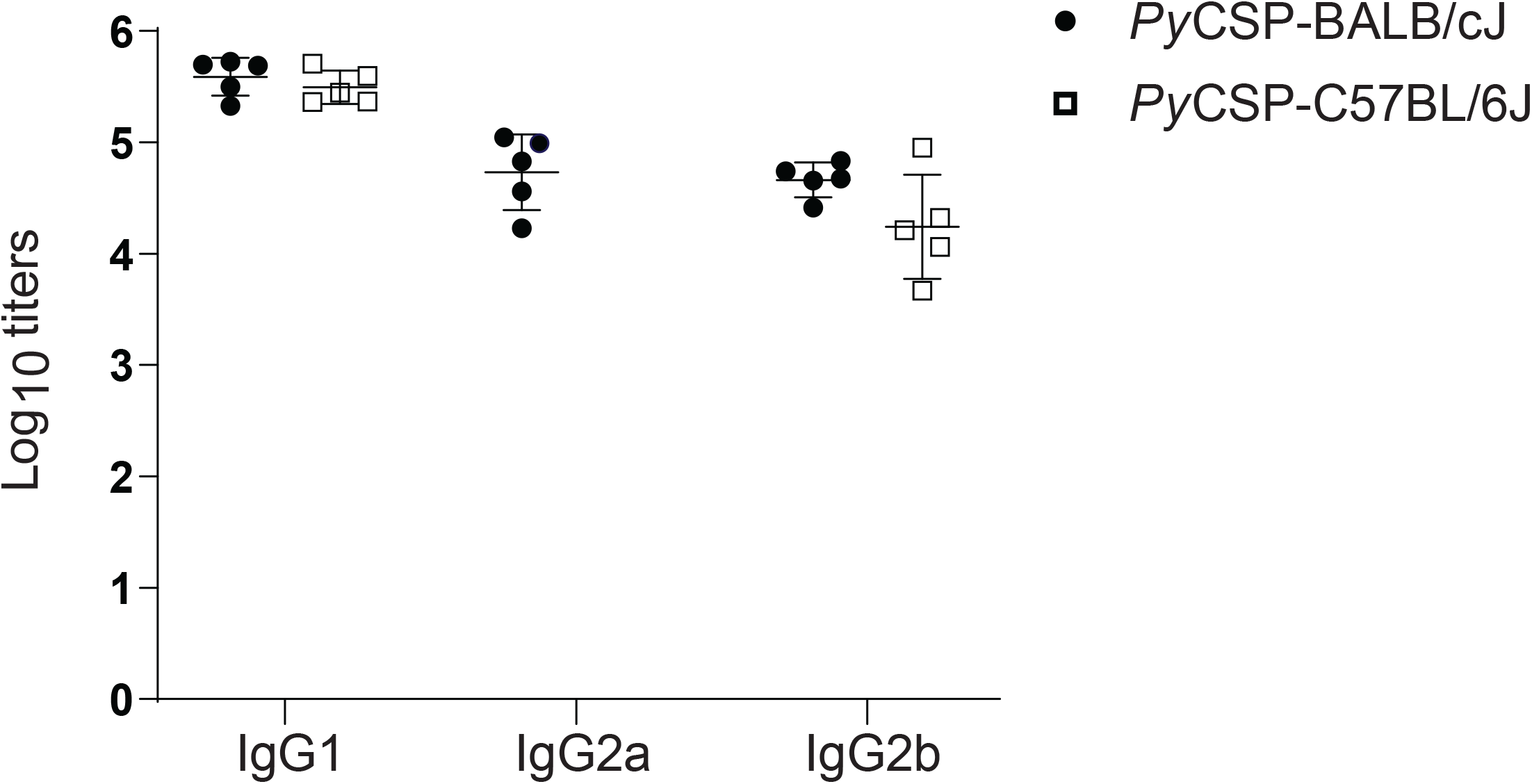
IgG subclass differences between the *Py*CSP-immunized BALB/cJ and C57BL/6J mice. The plasma IgG1, IgG2a (only for BALB/cJ) and IgG2b antibody responses of *Py*CSP-immunized BALB/cJ (black circles) and C57BL/6J (black squares) mice at week 7 (pre-challenge) were measured by ELISA using biotinylated *Py*CSP as ligand. Data analyzed by Two-way ANOVA and p values were obtained by Sidak’s multiple comparisons test. *\*\*\*\*p<0*.*0001*

**Figure S3.**
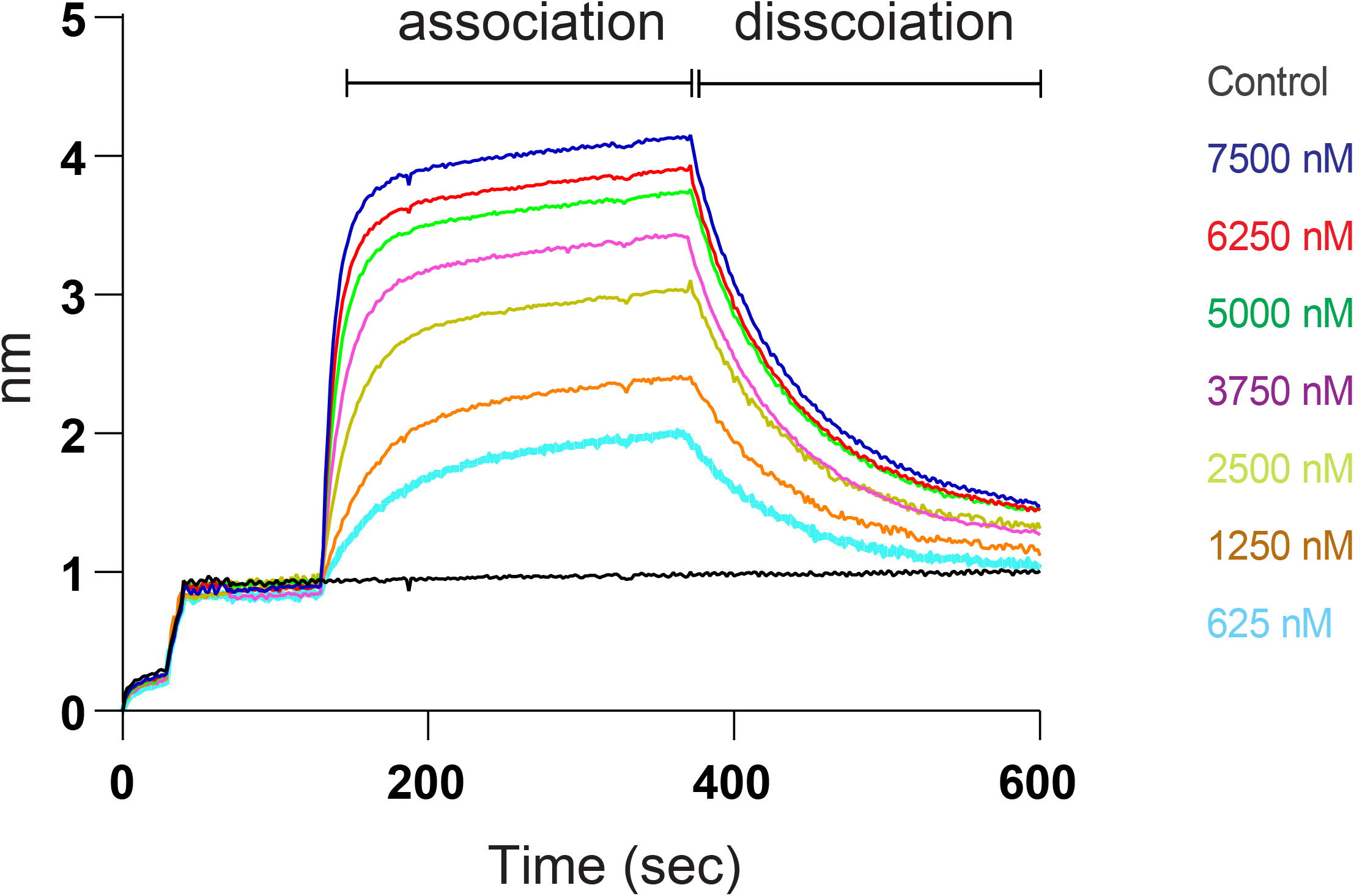
RAM2 Fab association and dissociation kinetics. Streptavidin biosensors loaded with *N*-terminally biotinylated-major repeat peptides were incubated in different concentrations of RAM2 Fab ranging from 7500 nM to 625 nM and the association and dissociation kinetics were analyzed. The resulting association and dissociation sensorgrams were analyzed by a global fit 1:1 binding model using the ForteBio data analysis software (version 7.0.1.5) generating K_D_ as estimated from the on- and off-rates.

**Figure S4.**
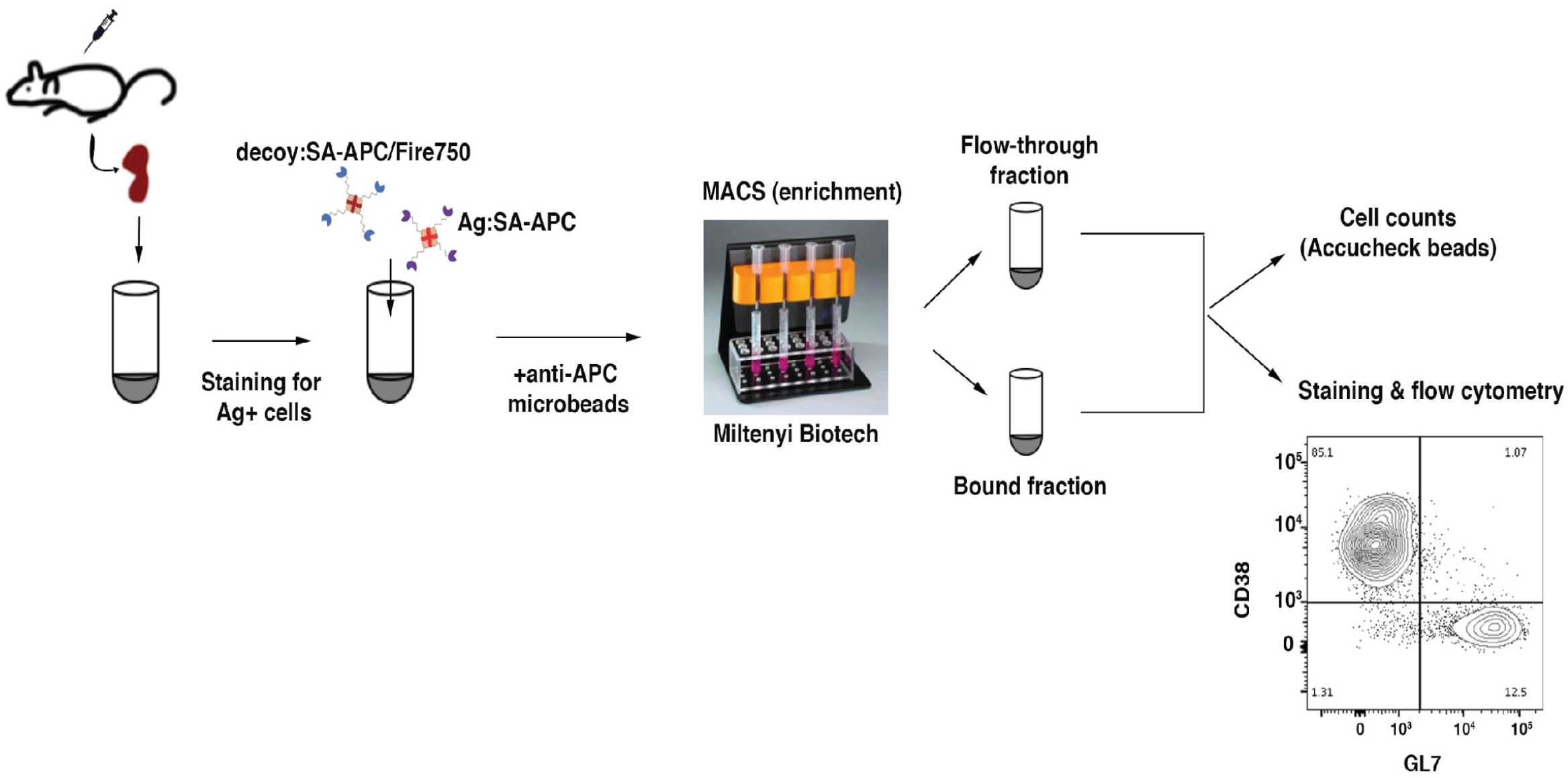
Summary of the workflow showing the different steps involved in the quantification of CSP-specific B-cell responses as described in the Materials and Methods section.

**Figure S5.**
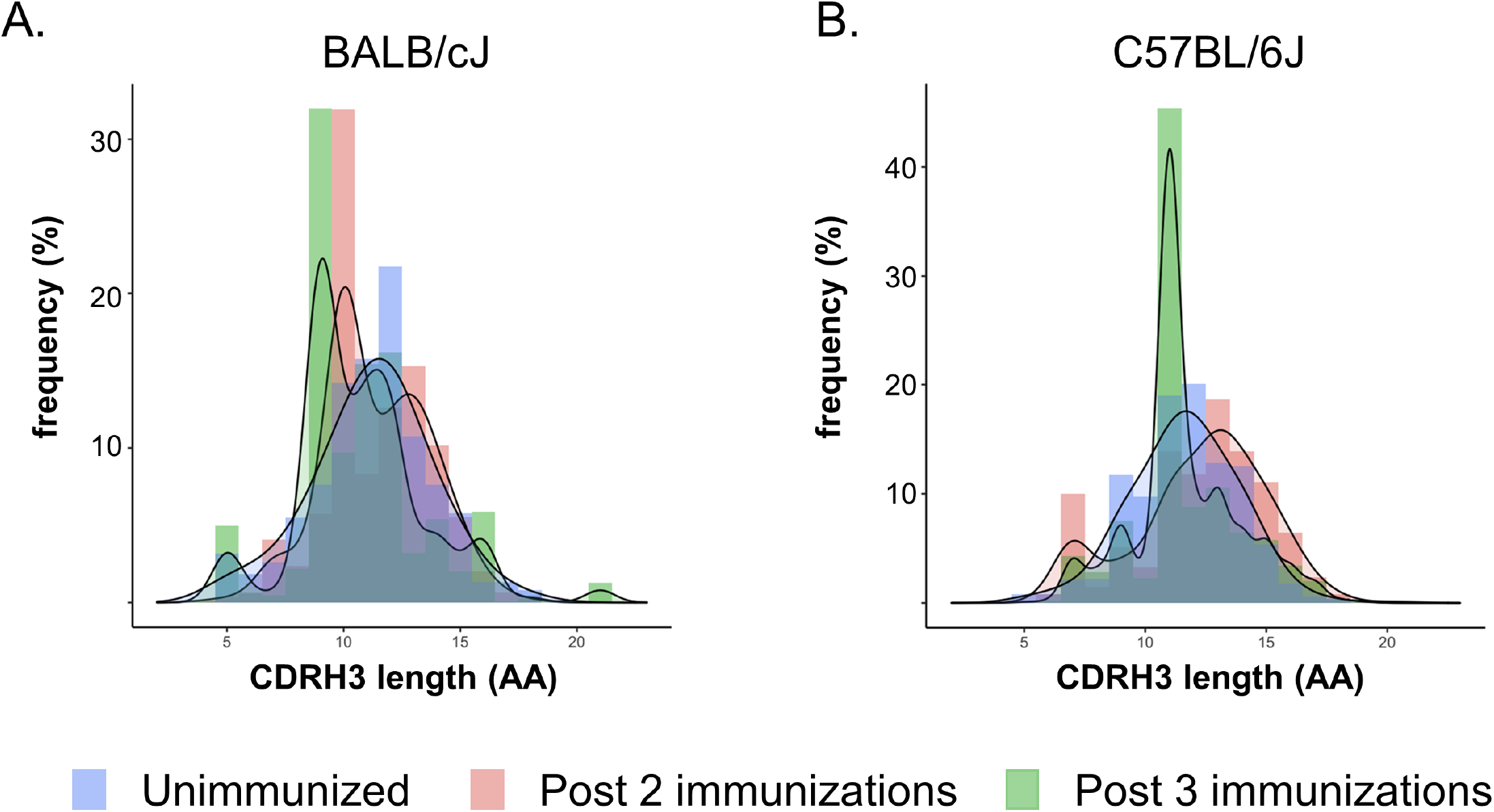
The CDRH3 amino acids (aa) lengths and density of **(A)** *Py*CSP-specific MBCs γ chain sequences from naïve animals (lavender), and after two (peach) or three (green) immunizations in BALB/cJ mice, and **(B)** from C57BL/6J mice. The results are representative of three independent experiments (n=5/group).

## References

Alexander, M.R., Ringe, R., Sanders, R.W., Voss, J.E., Moore, J.P., and Klasse, P.J. (2015). What Do Chaotrope-Based Avidity Assays for Antibodies to HIV-1 Envelope Glycoproteins Measure? J Virol 89, 5981–5995.

Aliprandini, E., Tavares, J., Panatieri, R.H., Thiberge, S., Yamamoto, M.M., Silvie, O., Ishino, T., Yuda, M., Dartevelle, S., Traincard, F., et al. (2018). Cytotoxic anti-circumsporozoite antibodies target malaria sporozoites in the host skin. Nat Microbiol 3, 1224–1233.

Balaban, A.E., Kanatani, S., Mitra, J., Gregory, J., Vartak, N., Sinnis-Bourozikas, A., Frischknecht, F., Ha, T., and Sinnis, P. (2021). The repeat region of the circumsporozoite protein is an elastic linear spring with a functional role in <em>Plasmodium</em> sporozoite motility. bioRxiv, 2021.2005.2012.443759.

Benhnini, F., Chenik, M., Laouini, D., Louzir, H., Cazenave, P.A., and Dellagi, K. (2009). Comparative evaluation of two vaccine candidates against experimental leishmaniasis due to Leishmania major infection in four inbred mouse strains. Clin Vaccine Immunol 16, 1529–1537.

Bruna-Romero, O., Rocha, C.D., Tsuji, M., and Gazzinelli, R.T. (2004). Enhanced protective immunity against malaria by vaccination with a recombinant adenovirus encoding the circumsporozoite protein of Plasmodium lacking the GPI-anchoring motif. Vaccine 22, 3575–3584.

Burrows, J.N., Duparc, S., Gutteridge, W.E., Hooft van Huijsduijnen, R., Kaszubska, W., Macintyre, F., Mazzuri, S., Mohrle, J.J., and Wells, T.N.C. (2017). New developments in anti-malarial target candidate and product profiles. Malar J 16, 26.

Camponovo, F., Campo, J.J., Le, T.Q., Oberai, A., Hung, C., Pablo, J.V., Teng, A.A., Liang, X., Sim, B.K.L., Jongo, S., et al. (2020). Proteome-wide analysis of a malaria vaccine study reveals personalized humoral immune profiles in Tanzanian adults. Elife 9.

Collins, K.A., Brod, F., Snaith, R., Ulaszewska, M., Longley, R.J., Salman, A.M., Gilbert, S.C., Spencer, A.J., Franco, D., Ballou, W.R., et al. (2021). Ultra-low dose immunization and multi-component vaccination strategies enhance protection against malaria in mice. Sci Rep 11, 10792.

Datoo, M.S., Natama, M.H., Some, A., Traore, O., Rouamba, T., Bellamy, D., Yameogo, P., Valia, D., Tegneri, M., Ouedraogo, F., et al. (2021). Efficacy of a low-dose candidate malaria vaccine, R21 in adjuvant Matrix-M, with seasonal administration to children in Burkina Faso: a randomised controlled trial. Lancet 397, 1809–1818.

Doepker, L.E., Simonich, C.A., Ralph, D., Shipley, M.M., Garrett, M., Gobillot, T., Vigdorovich, V., Sather, D.N., Nduati, R., Matsen, F.A.t., et al. (2020). Diversity and Function of Maternal HIV-1-Specific Antibodies at the Time of Vertical Transmission. J Virol 94.

Douglass, A.N., Metzger, P.G., Kappe, S.H.I., and Kaushansky, A. (2015). Flow Cytometry-Based Assessment of Antibody Function Against Malaria Pre-erythrocytic Infection. In Malaria Vaccines: Methods and Protocols, A. Vaughan, ed. (New York, NY: Springer New York), pp. 49–58.

Epstein, J.E., Tewari, K., Lyke, K.E., Sim, B.K., Billingsley, P.F., Laurens, M.B., Gunasekera, A., Chakravarty, S., James, E.R., Sedegah, M., et al. (2011). Live attenuated malaria vaccine designed to protect through hepatic CD8(+) T cell immunity. Science 334, 475–480.

Ewer, K.J., O’Hara, G.A., Duncan, C.J., Collins, K.A., Sheehy, S.H., Reyes-Sandoval, A., Goodman, A.L., Edwards, N.J., Elias, S.C., Halstead, F.D., et al. (2013). Protective CD8+ T-cell immunity to human malaria induced by chimpanzee adenovirus-MVA immunisation. Nat Commun 4, 2836.

Ewer, K.J., Sierra-Davidson, K., Salman, A.M., Illingworth, J.J., Draper, S.J., Biswas, S., and Hill, A.V. (2015). Progress with viral vectored malaria vaccines: A multi-stage approach involving “unnatural immunity”. Vaccine 33, 7444–7451.

Ghosh, A.K., and Jacobs-Lorena, M. (2009). Plasmodium sporozoite invasion of the mosquito salivary gland. Curr Opin Microbiol 12, 394–400.

Giudicelli, V., Chaume, D., and Lefranc, M.P. (2005). IMGT/GENE-DB: a comprehensive database for human and mouse immunoglobulin and T cell receptor genes. Nucleic Acids Res 33, D256–261.

Jongo, S.A., Shekalaghe, S.A., Church, L.W.P., Ruben, A.J., Schindler, T., Zenklusen, I., Rutishauser, T., Rothen, J., Tumbo, A., Mkindi, C., et al. (2018). Safety, Immunogenicity, and Protective Efficacy against Controlled Human Malaria Infection of Plasmodium falciparum Sporozoite Vaccine in Tanzanian Adults. Am J Trop Med Hyg 99, 338–349.

Kisalu, N.K., Idris, A.H., Weidle, C., Flores-Garcia, Y., Flynn, B.J., Sack, B.K., Murphy, S., Schön, A., Freire, E., Francica, J.R., et al. (2018). A human monoclonal antibody prevents malaria infection by targeting a new site of vulnerability on the parasite. Nat Med 24, 408–416.

Kuipers, K., van Selm, S., van Opzeeland, F., Langereis, J.D., Verhagen, L.M., Diavatopoulos, D.A., and de Jonge, M.I. (2017). Genetic background impacts vaccine-induced reduction of pneumococcal colonization. Vaccine 35, 5235–5241.

Miller, J.L., Sack, B.K., Baldwin, M., Vaughan, A.M., and Kappe, S.H.I. (2014). Interferon-mediated innate immune responses against malaria parasite liver stages. Cell Rep 7, 436–447.

Mordmuller, B., Surat, G., Lagler, H., Chakravarty, S., Ishizuka, A.S., Lalremruata, A., Gmeiner, M., Campo, J.J., Esen, M., Ruben, A.J., et al. (2017). Sterile protection against human malaria by chemoattenuated PfSPZ vaccine. Nature 542, 445–449.

Murugan, R., Buchauer, L., Triller, G., Kreschel, C., Costa, G., Pidelaserra Martí, G., Imkeller, K., Busse, C.E., Chakravarty, S., Sim, B.K.L., et al. (2018). Clonal selection drives protective memory B cell responses in controlled human malaria infection. Sci Immunol 3.

Olotu, A., Fegan, G., Wambua, J., Nyangweso, G., Leach, A., Lievens, M., Kaslow, D.C., Njuguna, P., Marsh, K., and Bejon, P. (2016). Seven-Year Efficacy of RTS,S/AS01 Malaria Vaccine among Young African Children. N Engl J Med 374, 2519–2529.

Pullen, G.R., Fitzgerald, M.G., and Hosking, C.S. (1986). Antibody avidity determination by ELISA using thiocyanate elution. J Immunol Methods 86, 83–87.

Rognes, T., Flouri, T., Nichols, B., Quince, C., and Mahe, F. (2016). VSEARCH: a versatile open source tool for metagenomics. PeerJ 4, e2584.

Sack, B.K., Miller, J.L., Vaughan, A.M., Douglass, A., Kaushansky, A., Mikolajczak, S., Coppi, A., Gonzalez-Aseguinolaza, G., Tsuji, M., Zavala, F., et al. (2014). Model for in vivo assessment of humoral protection against malaria sporozoite challenge by passive transfer of monoclonal antibodies and immune serum. Infect Immun 82, 808–817.

Scally, S.W., Murugan, R., Bosch, A., Triller, G., Costa, G., Mordmuller, B., Kremsner, P.G., Sim, B.K.L., Hoffman, S.L., Levashina, E.A., et al. (2018). Rare PfCSP C-terminal antibodies induced by live sporozoite vaccination are ineffective against malaria infection. J Exp Med 215, 63–75.

Shugay, M., Britanova, O.V., Merzlyak, E.M., Turchaninova, M.A., Mamedov, I.Z., Tuganbaev, T.R., Bolotin, D.A., Staroverov, D.B., Putintseva, E.V., Plevova, K., et al. (2014). Towards error-free profiling of immune repertoires. Nat Methods 11, 653–655.

Simonich, C.A., Doepker, L., Ralph, D., Williams, J.A., Dhar, A., Yaffe, Z., Gentles, L., Small, C.T., Oliver, B., Vigdorovich, V., et al. (2019). Kappa chain maturation helps drive rapid development of an infant HIV-1 broadly neutralizing antibody lineage. Nat Commun 10, 2190.

Sissoko, M.S., Healy, S.A., Katile, A., Omaswa, F., Zaidi, I., Gabriel, E.E., Kamate, B., Samake, Y., Guindo, M.A., Dolo, A., et al. (2017). Safety and efficacy of PfSPZ Vaccine against Plasmodium falciparum via direct venous inoculation in healthy malaria-exposed adults in Mali: a randomised, double-blind phase 1 trial. Lancet Infect Dis 17, 498–509.

Stoute, J.A., Slaoui, M., Heppner, D.G., Momin, P., Kester, K.E., Desmons, P., Wellde, B.T., Garcon, N., Krzych, U., and Marchand, M. (1997). A preliminary evaluation of a recombinant circumsporozoite protein vaccine against Plasmodium falciparum malaria. RTS,S Malaria Vaccine Evaluation Group. N Engl J Med 336, 86–91.

Sun, P., Schwenk, R., White, K., Stoute, J.A., Cohen, J., Ballou, W.R., Voss, G., Kester, K.E., Heppner, D.G., and Krzych, U. (2003). Protective immunity induced with malaria vaccine, RTS,S, is linked to Plasmodium falciparum circumsporozoite protein-specific CD4+ and CD8+ T cells producing IFN-gamma. J Immunol 171, 6961–6967.

Tan, J., Sack, B.K., Oyen, D., Zenklusen, I., Piccoli, L., Barbieri, S., Foglierini, M., Fregni, C.S., Marcandalli, J., Jongo, S., et al. (2018). A public antibody lineage that potently inhibits malaria infection through dual binding to the circumsporozoite protein. Nat Med 24, 401–407.

Thai, E., Costa, G., Weyrich, A., Murugan, R., Oyen, D., Flores-Garcia, Y., Prieto, K., Bosch, A., Valleriani, A., Wu, N.C., et al. (2020). A high-affinity antibody against the CSP N-terminal domain lacks Plasmodium falciparum inhibitory activity. J Exp Med 217.

Triller, G., Scally, S.W., Costa, G., Pissarev, M., Kreschel, C., Bosch, A., Marois, E., Sack, B.K., Murugan, R., Salman, A.M., et al. (2017). Natural Parasite Exposure Induces Protective Human Anti-Malarial Antibodies. Immunity 47, 1197–1209 e1110.

Turchaninova, M.A., Davydov, A., Britanova, O.V., Shugay, M., Bikos, V., Egorov, E.S., Kirgizova, V.I., Merzlyak, E.M., Staroverov, D.B., Bolotin, D.A., et al. (2016). High-quality full-length immunoglobulin profiling with unique molecular barcoding. Nat Protoc 11, 1599–1616.

Vanderberg, J., Nussenzweig, R., and Most, H. (1969). Protective immunity produced by the injection of x-irradiated sporozoites of Plasmodium berghei. V. In vitro effects of immune serum on sporozoites. Mil Med 134, 1183–1190.

Vigdorovich, V., Oliver, B.G., Carbonetti, S., Dambrauskas, N., Lange, M.D., Yacoob, C., Leahy, W., Callahan, J., Stamatatos, L., and Sather, D.N. (2016). Repertoire comparison of the B-cell receptor-encoding loci in humans and rhesus macaques by next-generation sequencing. Clin Transl Immunology 5, e93.

Vijayan, K., Visweswaran, G.R.R., Chandrasekaran, R., Trakhimets, O., Brown, S.L., Watson, A., Zuck, M., Dambrauskas, N., Raappana, A., Carbonetti, S., et al. (2021). Antibody interference by a non-neutralizing antibody abrogates humoral protection against Plasmodium yoelii liver stage. Cell Rep 36, 109489.

WHO (2019). World malaria report 2019. World Health Organization, Geneva.

WHO (2020). World malaria report 2020: 20 years of global progress and challenges. In World Health Organization, Geneva.

Ye, J., Ma, N., Madden, T.L., and Ostell, J.M. (2013). IgBLAST: an immunoglobulin variable domain sequence analysis tool. Nucleic Acids Res 41, W34–40.

